# MORC-1 is a key component of the *C. elegans* CSR-1 germline gene licensing mechanism

**DOI:** 10.1101/2024.10.02.616347

**Authors:** Jessica A. Kirshner, Colette L. Picard, Natasha E. Weiser, Nicita Mehta, Suhua Feng, Victoria N. Murphy, Anna Vakhnovetsky, Amelia F. Alessi, Connie Xiao, Kai Inoki, Sonia El Mouridi, Christian Frøkjær-Jensen, Steven E. Jacobsen, John K. Kim

## Abstract

The Argonaute CSR-1 is essential for germline development in *C. elegans*. Mutation of *csr-1* downregulates thousands of germline-expressed genes, leading to the model that the CSR-1-mediated small RNA pathway promotes, or “licenses,” gene expression by an unknown mechanism. CSR-1 also silences a limited number of genes through its canonical endonucleolytic “slicer” activity. We show that the GHKL-type ATPase MORC-1, a CSR-1 slicing target, over-accumulates at CSR-1 “licensed” target genes in *csr-1*(*-*), which correlates with ectopic gain of H3K9me3, H3K36me3 loss, and gene downregulation. Loss of *morc-1* rescues *csr-1*(*-*) defects, while overexpressing MORC-1 in the germline of wild-type worms is sufficient to cause sterility and downregulate CSR-1 targets. These results show that MORC-1 overexpression in *csr-1*(*-*) is a primary driver of the CSR-1-mediated gene licensing mechanism.

**One-Sentence Summary:** MORC-1 acts downstream of CSR-1 to regulate germline chromatin states and is a key component of the gene licensing mechanism.

## Main Text

Maintaining genome integrity is particularly important in the germline. Germline gene expression in *C. elegans* is tightly regulated by a series of small RNA pathways working coordinately to ensure that germline genes are expressed, while non-germline genes and foreign genetic elements are stably silenced. Small RNA-dependent surveillance and silencing of transposons and other deleterious genetic elements is mediated by proteins in the highly conserved Argonaute family (*1*, *2*). However, the Argonaute CSR-1 associates with a distinct set of endo-siRNAs (22G-RNAs) complementary to ∼4000 endogenous protein-coding genes and somehow promotes, or “licenses,” their expression (*3*). Consequently, loss of *csr-1* causes modest downregulation of most CSR-1 “licensed” target genes (*3*, *4*), which constitute the majority of the genes expressed in the germline. This gene licensing pathway is essential for proper germline development, as *csr-1*(-) mutant worms are completely sterile (*2*, *3*). However, despite further characterization (*5–8*), the mechanism by which CSR-1 promotes the expression of its target genes remains largely unknown, and the sterility phenotype of *csr-1*(*-*) worms has complicated study of this pathway.

In addition to its gene licensing function, CSR-1 has endonucleolytic “slicing” activity (*9*, *10*), like many other Argonaute proteins. About 100 of the ∼4000 CSR-1 targets are cleaved and silenced, rather than licensed (*10*). One of the reported CSR-1 silenced targets is *morc-1*, the sole *C. elegans* homolog of the conserved family of *Microrchidia* GHKL-type ATPases (*10*). MORCs are highly conserved in plants and animals, with diverse functions including repressing transcription and maintaining repressive chromatin states (*11–19*). The specific mechanisms by which MORCs repress transcription vary. Some mammalian MORCs are thought to regulate H3K9me3 via interactions with the SETDB1/HUSH complex (*14*, *19*), while several *Arabidopsis* MORCs promote the establishment of DNA methylation (*18*). *C. elegans* MORC-1 maintains silencing at some sites by preventing active chromatin from spreading into repressive regions marked by H3K9me3 (*15*), and, *in vitro*, can bind and compact chromatin independent of DNA sequence (*16*). However, the *in vivo* targets and function of *C. elegans* MORC-1 remain largely uncharacterized. Additionally, like many MORCs in other organisms, *C. elegans* MORC-1 is important for proper germline development (*15*). Here, we demonstrate that targeted silencing of *morc-1* by CSR-1 is a major cause of the CSR-1-mediated germline gene licensing mechanism, and that MORC-1 overexpression alone recapitulates *csr-1* mutant phenotypes, including CSR-1 target downregulation and sterility.

### *morc-1*(-) partially rescues *csr-1* sterility

*C. elegans* lacking *csr-1* have a defective germline and are sterile, and the few embryos that do form die by the 100-cell stage (*3*). We hypothesized that this defect could be due to the overexpression of one or more CSR-1 cleavage targets. We examined the fertility of wild-type and *morc-1*(-) worms on either control (empty vector, EV) or *csr-1* RNAi, and found that while wild-type worms grown on *csr-1* RNAi are fully sterile, *morc-1*(-) worms on *csr-1* RNAi retain partial fertility (Fig. 1A). To further characterize the nature of the interaction between *morc-1* and *csr-1*, we used CRISPR/Cas9 to generate two novel, viable *csr-1* mutants (Fig 1B). The first, *csr-1 (G560R)*, contains a point mutation homologous to the anti-morph mutation in Argonaute ALG-1 that interferes with microRNA passenger strand removal (*20*), and is predicted to interfere with small RNA binding. The second, *aid::csr-1*, contains an auxin-inducible degron (AID) tag, allowing CSR-1 to be degraded in the presence of the plant hormone auxin and the F-box protein TIR1 (*21*), which we express under a germline-specific promoter (Fig 1B, fig. S1A). These mutants affect both *csr-1* isoforms, *csr-1a* and *csr-1b.* (Fig. 1B). To confirm that both strains conditionally act as true *csr-1* mutants, we assessed their fertility and gene expression profiles at elevated temperature (*csr-1(G560R)*) or on an auxin concentration gradient (*aid::csr-1*). Fertility was abolished in each line (fig. S1B). Both novel mutants additionally displayed the CSR-1 gene licensing phenotype, characterized by mild downregulation of CSR-1 licensed targets (*4*), while CSR-1 slicing targets (*10*) were upregulated as expected (fig. S1C). CSR-1-bound 22G-RNAs were also moderately destabilized in both mutants, but particularly in *aid::csr-1*, likely due to *csr-1 (G560R)* retaining residual ability to bind small RNAs (fig. S1D). We conclude that both *csr-1(G560R)* and *aid::csr-1* can conditionally phenocopy *csr-1*(*-*).

**Fig. 1.**
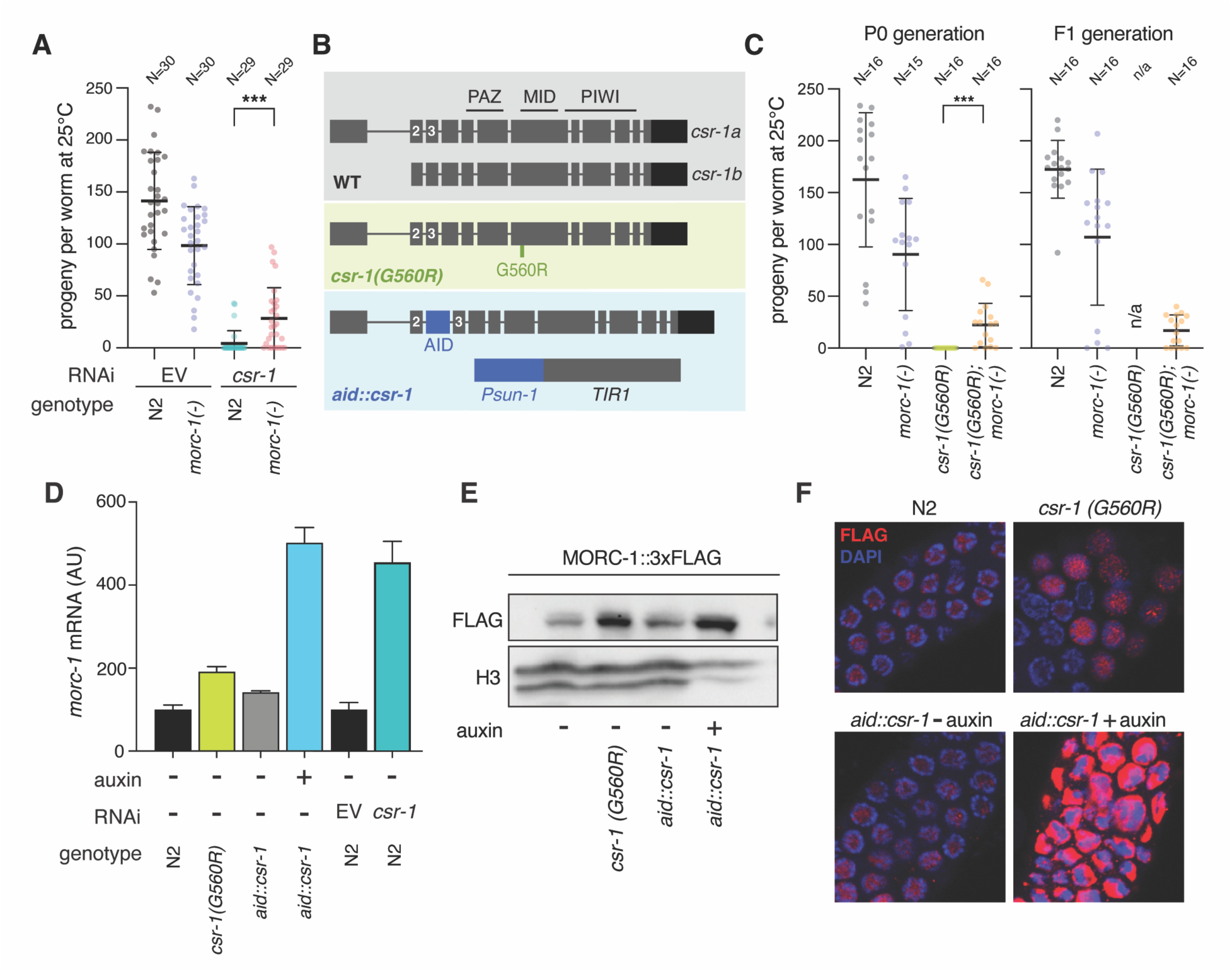
*morc-1*(*-*) is a suppressor of *csr-1.* (**A**) Fertility of wild-type (N2) or *morc-1*(*-*) worms grown on either empty vector (EV) or *csr-1* RNAi. Each point represents the progeny produced by an individual worm. (**B**) Diagram of CSR-1 protein structure for both isoforms, as well as the two conditional *csr-1* mutants generated in this study: *aid::csr-1* and *csr-1(G560R)*. (**C**) Fertility of wild-type, *morc-1*(*-*)*, csr-1(G560R)* and *csr-1(G560R); morc-1*(*-*) double mutant worms. Each point represents the progeny produced by an individual worm. Left panel shows first generation (P0) grown at the *csr-1(G560R)* non-permissive temperature of 25°C, while the right panel shows fertility of second generation (F1), also grown at 25°C. Because *csr-1(G560R)* P0 worms did not produce any progeny at 25°C, their fertility in the F1 generation could not be assayed. (**D**) Upregulation of *morc-1* mRNA in both *csr-1(G560R)* and *aid::csr-1*, as well as on *csr-1* RNAi, by qPCR. Error bars represent standard deviation between two technical replicates. (**E**) Western blot of MORC-1::3xFLAG protein in both *csr-1(G560R)* and *aid::csr-1*, with H3 as a loading control. (**F**) Immunofluorescence of MORC-1::3xFLAG (red) in *csr-1(G560R)* and *aid::csr-1* in dissected germlines of indicated genotype and treatment. (D-F) Worms were treated with either 0 μM [(-) auxin] or 100 μM auxin [(+) auxin]. (A,B) *** = p < 0.001, one-tailed t-test.

We next tested the fertility of *csr-1 (G560R)* relative to *csr-1 (G560R); morc-1*(-) and confirmed that *morc-1*(-) also partially rescues the sterility defect of *csr-1 (G560R)* (Fig 1C, fig. S2). Fertility in the double mutant was maintained transgenerationally (Fig. 1C), highlighting the stability of this rescue and confirming that *morc-1* is a suppressor of *csr-1*.

### MORC-1 is overexpressed in *csr-1* mutants

We next investigated the molecular mechanism of the rescue of *csr-1* by *morc-1*(*-*). Gerson-Gurwitz *et al.* 2016 (*10*) identified 133 putative slicing targets of CSR-1 that become upregulated in a slicing inactive *csr-1* mutant, one of which was *morc-1*. To confirm that CSR-1 represses *morc-1*, we investigated *morc-1* expression using our new *csr-1* mutants. We found that *morc-1* was upregulated at both the mRNA and protein levels compared to wild-type worms, both in our *csr-1* mutant strains and in wild-type worms treated with *csr-1* RNAi (Fig. 1D-F, fig. S3A-D). This increase in *morc-1* RNA and protein was specific to *csr-1,* as *morc-1* was not upregulated upon knockdown of other CSR-1 pathway components (fig. S3B-D). This suggests that *morc-1* is directly cleaved by CSR-1, rather than being regulated by the broader CSR-1 pathway.

### MORC-1 binds the transcriptional start sites of germline-expressed genes and spreads in *csr-1* mutants

*C. elegans* MORC-1 is a highly conserved DNA binding protein that can condense DNA and chromatin *in vitro* (*16*), but its function *in vivo* is not yet understood. To investigate the consequences of MORC-1 overexpression in a *csr-1* mutant background, we first examined the chromatin binding profile of MORC-1 in wild-type worms. We performed ChIP-seq in purified germline nuclei derived from a strain expressing MORC-1 tagged with a 3xFlag epitope at its endogenous locus by CRISPR/Cas9 ((*22*), see methods). Despite the conserved role of MORCs in silencing, *C. elegans* MORC-1 was depleted from transposable elements (TEs) and repeats in the germline (Fig. 2A,B, fig. S4A). Instead, MORC-1 bound specifically at the transcriptional start sites (TSSs) of protein coding genes (Fig. 2A,B, fig. S4A-D). MORC-1 binding correlated well with germline gene expression (fig. S4B,D). Correspondingly, CSR-1 targets, which tend to be highly expressed in the germline (fig. S5A), as well as other germline-expressed genes (*23*), were strongly bound by MORC-1 (Fig. 2B,C, fig. S5B-C). The genes with the highest MORC-1 levels in wild-type animals, including CSR-1 targets, show mild upregulation in a published *morc-1*(*-*) RNA-seq dataset (*15*) (fig. S6A-C). This suggests that despite binding primarily to the promoters of highly expressed genes, MORC-1 binding may nonetheless have a repressive effect.

**Fig. 2.**
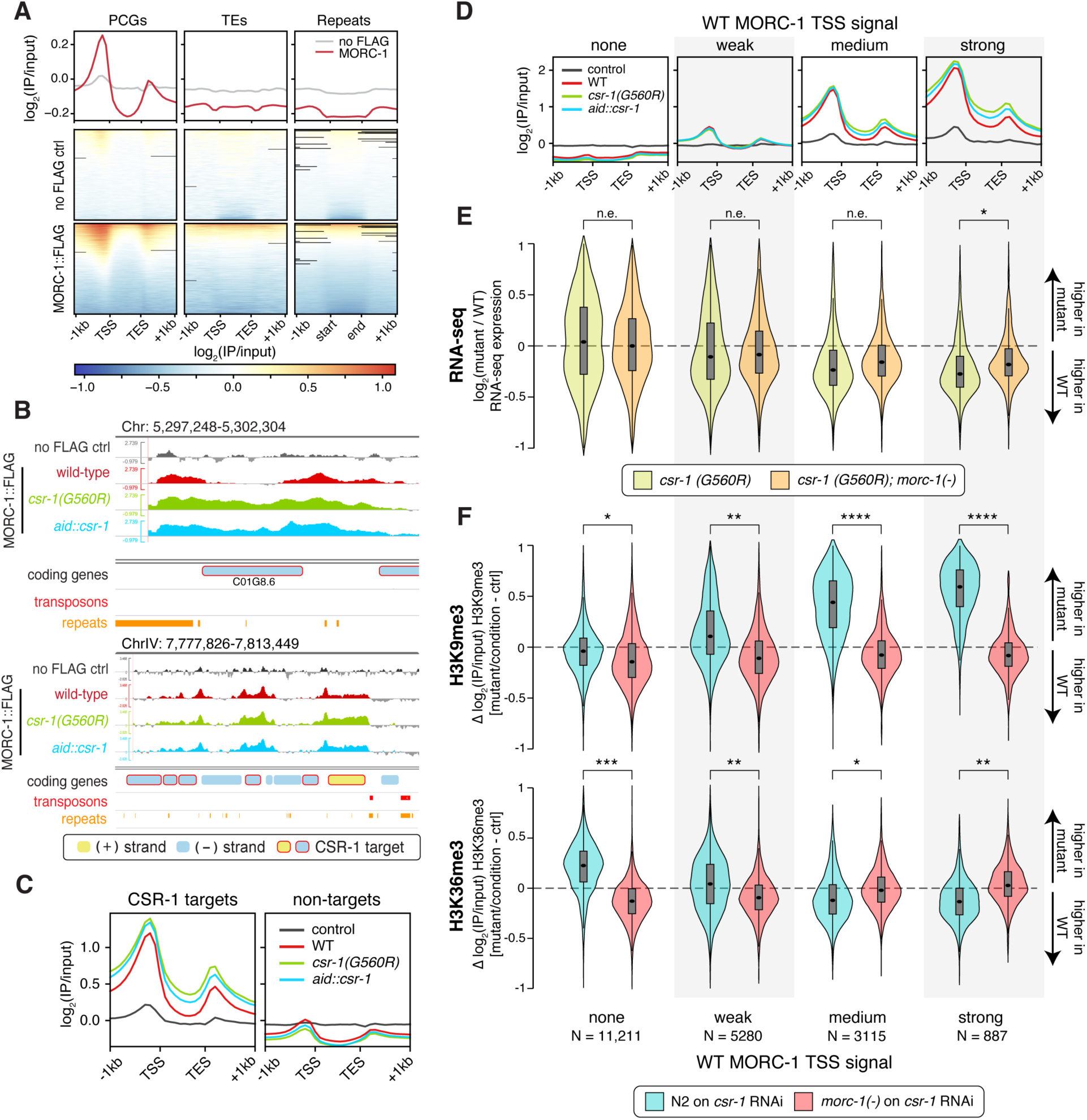
Chromatin and expression changes in *csr-1* mutants correlate with MORC-1 overaccumulation and are rescued by *morc-1*(*-*). (**A**) Metaplots and heatmaps of anti-FLAG ChIP-seq signal from purified germline nuclei of the MORC-1::3xFLAG expressing strain or control wild-type worms lacking FLAG. ChIP-seq signal over protein-coding genes (PCGs), transposons (TEs) and repeat regions (repeats) are shown. Features were scaled to 1kb length. Each IP sample was normalized to a matched input sample, and the log of that ratio is plotted (log_2_(IP/input)), average of two replicates. (**B**) Example browser images showing average log_2_(IP/input) MORC-1::3xFLAG signal in wild-type germline (red), *csr-1(G560R)* (green) and *aid::csr-1* (blue), as well as the no-FLAG control (grey). Locations of genes, transposons, and repeats shown on bottom tracks. Genes on the forward and reverse strands are colored yellow and blue, respectively, while CSR-1 targets are circled red. (**C-D**) Average germline log_2_(IP/input) MORC-1::3xFLAG signal over protein-coding genes in wild-type (red), *csr-1(G560R)* (green), *aid::csr-1* (blue), and a control sample lacking FLAG (grey). Genes were scaled to 1kb length. Plots were made over (C) CSR-1 targets vs. non-targets, and (D) genes binned by promoter MORC-1 levels (see Data S2). (**E**) Change in gene expression in indicated mutant vs. wild-type control, across genes binned into four different groups based on wild-type levels of MORC-1 at their transcriptional start sites (TSS). Gene bins same as in (D). Expression change shown is the log2(fold change) value estimated by DESeq2 (*35*). Number of genes in each bin shown at the bottom of panel (F). (**F**) Difference in H3K9me3 and H3K36me3 ChIP-seq signal (log_2_(IP/input)) in indicated mutant and/or condition, compared to wild-type worms treated with control RNAi. Genes were again binned based on MORC-1 levels at the TSS in wild-type, as in (D) and (E). (E-F) Stars indicate effect size as measured using Cohen’s d. n.e. = no/minimal effect (|d| < 0.2), * = |d| > 0.2, ** = |d| > 0.5, *** = |d| > 0.9, **** = |d| > 1.5. All comparisons give p ∼ 0 by Student’s t-test due to large sample size.

We then performed ChIP-seq for MORC-1::3xFlag in our novel *csr-1* mutant strains, where MORC-1 becomes overexpressed (Fig. 1D-F). While MORC-1 was restricted to TSSs in wild-type, in both *csr-1* mutants, its binding increased further at the promoters of genes and spread into the gene bodies of its targets, where it is normally depleted (Fig. 2B-D, fig. S7A-C). MORC-1 accumulation, when overexpressed, occurred most prominently at targets already abundantly bound by MORC-1 in wild-type, and did not occur at non-targets (fig. S7A-C). Therefore, CSR-1 targets and other genes strongly bound by MORC-1 in wild-type became further enriched for MORC-1 in *csr-1* mutants, while non-targets, repeats, and transposable elements remained depleted for MORC-1 binding (Fig. 2B-D, fig. S7A-C).

### MORC-1 spreading in *csr-1* represses gene expression

We hypothesized that the spread of MORC-1 across CSR-1 targets might contribute to the downregulation of these genes in *csr-1* mutants. Indeed, the genes that gain the most MORC-1 in *csr-1(G560R)* and *aid::csr-1* were significantly downregulated in both strains, while genes with little or no MORC-1 were not affected (Fig. 2E, fig. S8A-C). To investigate this further, we clustered genes based on the change in MORC-1 levels in our two *csr-1* mutants using *k-*means clustering (fig. S9A). Both *csr-1* mutants showed very similar patterns. One cluster of genes strongly gained MORC-1 over the entire gene body and promoter in *csr-1*, while others only gained MORC-1 in the 5’ region, or in the 3’ region (fig. S9A). Other clusters exhibited either minimal MORC-1 gain or loss of MORC-1 in *csr-1*. The genes in clusters that strongly gained MORC-1 over either their entire length or their 5’ region showed the strongest downregulation in both *csr-1(G560R)* and *aid::csr-1* (clusters C1 and C2, fig. S9B). More generally, regression analysis revealed a modest but consistent correlation between gain of MORC-1 and gene downregulation in both *csr-1* strains (Fig. S9C). The clusters which showed strong MORC-1 gain in *csr-1* were predominantly CSR-1 targets and/or highly expressed in the germline (fig. S10A-C). These data suggest that ectopic MORC-1 gain over the promoter and/or gene body in *csr-1* mutants leads to gene downregulation, particularly at germline expressed genes and CSR-1 targets.

### *morc-1*(*-*) rescues *csr-1* defects in gene expression and chromatin states

If MORC-1 over-accumulation and spreading at CSR-1 target genes and other germline-expressed genes is responsible for their downregulation in *csr-1*, then loss of *morc-1* in *csr-1* should rescue this phenotype. We therefore performed RNA-seq in *csr-1(G560R); morc-1*(*-*). As expected, the downregulation of genes that strongly gained MORC-1 in *csr-1*, including CSR-1 targets, was partially rescued by loss of *morc-1*, while genes not bound by MORC-1 were not affected (Fig. 2E, fig. S11A,B). Expression of CSR-1 slicing targets was not rescued by loss of *morc-1*, consistent with these being regulated post-transcriptionally by direct CSR-1 slicing (fig. S11B). In fact, the slicing targets were even further upregulated in *csr-1(G560R); morc-1*(*-*). CSR-1 slicing targets gained substantial MORC-1 binding in our *csr-1* mutants (fig. S11C). Therefore, in addition to becoming upregulated post-transcriptionally by loss of CSR-1 slicing in *csr-1(G560R); morc-1*(*-*), the CSR-1 slicing targets are likely further upregulated due to loss of MORC-1-mediated transcriptional repression. Overall, these results show that loss of *morc-1* partially rescues the gene expression defects in *csr-1 (G560R)* at CSR-1 licensed targets and other genes strongly bound by MORC-1, suggesting that MORC-1 accumulation at affected genes in *csr-1* mutants is directly responsible for these expression defects.

*csr-1*(*-*) mutants have altered chromatin states (*4*, *24*). We therefore asked whether MORC-1 overexpression might be responsible for these changes. Loss of *morc-1* alone caused only mild loss of H3K9me3 and gain of H3K36me3 compared to wild-type (fig. S12A), as previously reported (*15*). Worms on *csr-1* RNAi instead had substantially increased H3K9me3 in over 2,000 1 kb bins genome-wide (fig. S12B). These bins were strongly depleted for H3K9me3 in wild-type worms, and highly enriched in protein-coding gene bodies (fig. S12C). Loss of *csr-1* also caused loss of H3K36me3 in approximately 900 1 kb bins, which were also disproportionally enriched in gene bodies and correlated with H3K9me3 gain (fig. S12B-C, S13A). These results suggest that H3K9me3, which marks repressive chromatin and is normally absent from gene bodies, ectopically spreads into genes when *csr-1* is depleted, while the active mark H3K36me3 is lost from these same genes, consistent with previous reports of disrupted chromatin boundaries in *csr-1* (*4*, *24*). We hypothesized that these changes in chromatin architecture in *csr-1* RNAi were a consequence of MORC-1 overaccumulation and spreading. Consistent with this, genes strongly bound by MORC-1 in wild-type, which also accumulate the most ectopic MORC-1 in *csr-1* mutants (Fig. 2D), showed the greatest gain of H3K9me3 and loss of H3K36me3 in *csr-1* RNAi (Fig. 2F, fig. S13A-C, fig. S14A-C). Additionally, *morc-1*(*-*) mutants on *csr-1* RNAi show complete rescue of this phenotype (Fig. 2F, fig. S13B-C, S14A-C, S15A). Conversely, *csr-1* RNAi does not rescue the mild chromatin defects of *morc-1* mutants (fig. S15B-C). These results indicate that ectopic MORC-1 gain over genes highly expressed in the germline, including CSR-1 targets, is responsible for the accumulation of H3K9me3 and loss of H3K36me3 at these genes in *csr-1* mutants, suggesting that MORC-1 regulates chromatin landscapes downstream of CSR-1.

### MORC-1 overexpression is sufficient to mimic *csr-1-*dependent germline gene licensing

Our data support a model in which MORC-1 normally binds to CSR-1 targets and other germline expressed genes, but over-accumulates at these sites in *csr-1*, leading to chromatin defects, gene downregulation, and sterility. To test this, we sought to overexpress MORC-1 in the germline using a transgene, to see whether this is sufficient to phenocopy *csr-1* mutant defects. This experiment is complicated, however, by the fact that MORC-1 promotes transgene silencing in *elegans* (*11*), so successful transgenic MORC-1 overexpression may cause silencing of the transgene. Additionally, if MORC-1 overexpression is lethal like *csr-1*, we would necessarily fail to recover worms overexpressing the transgene. Indeed, multiple independent approaches to generate MORC-1 overexpressing transgenic lines, using various genetic engineering strategies, were initially unsuccessful (Table S1).

We were ultimately able to generate a strain, *morcOE*, that conditionally overexpresses *morc-1* in the germline (see methods). To do this, we utilized a modular safe-harbor transgene insertion site (MosTi) (*25*) to integrate a single copy transgene containing codon-optimized *morc-1::3xflag* driven by a germline-specific promoter, into the *unc-119* locus (Fig. 3A). Also encoded on this transgene was a neomycin resistance gene and a neuronal mCherry marker. To circumvent lethality associated with *morc-1* overexpression, the codon-optimized *morc-1* was silenced via piRNAi utilizing an artificial piRNA expressed from an extrachromosomal array (*26*). The extrachromosomal piRNA array also contains a hygromycin resistance gene as a selectable marker. To induce *morc-1* overexpression conditionally, worms were removed from hygromycin to allow for the loss of the extrachromosomal piRNA array, and therefore loss of piRNAi-based silencing of codon-optimized *morc-1* (Fig. 3B). Consistent with *morc-1* overexpression being toxic and causing transgene silencing, we observed a high rate of silencing of the integrated *morc-1* transgene following loss of the extrachromosomal *morc-1*-silencing piRNA array. Worms that silenced the integrated *morc-1* transgene exhibited loss of neomycin resistance, loss of neuronal mCherry, and an uncoordinated (Unc) phenotype due to silencing at the *unc-119* locus where the *morc-1* transgene was integrated (Fig. 3B).

**Fig. 3.**
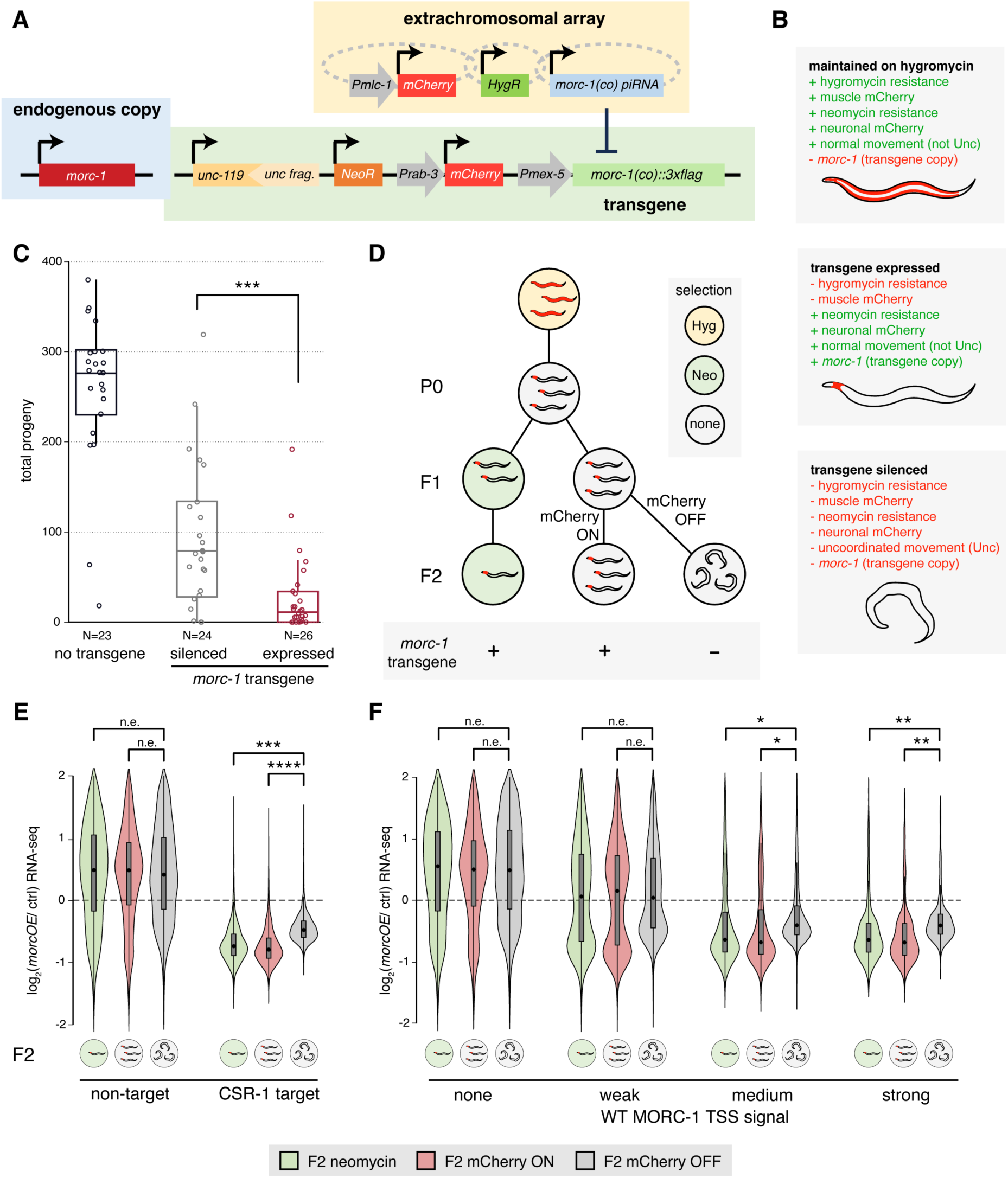
MORC-1 overexpression in wild-type germline phenocopies *csr-1* fertility and expression defects. (**A**) Schematic of our conditional germline MORC-1 overexpression line (*morcOE*, see methods). (**B**) Expected phenotypes of *morcOE* stock worms maintained on hygromycin (extrachromosomal array retained), worms removed from hygromycin that have lost the extrachromosomal array and express all genes on the integrated transgene, and worms that have silenced the integrated transgene. (**C**) Fertility of F2 *morcOE* worms not grown on neomycin, comparing worms that silenced the integrated *morc-1* transgene vs. those that did not. The transgene was considered silenced if either mCherry expression was lost or worms became uncoordinated (Unc; see B, Data S4). *** = p < 0.001, two-tailed t-test. (**D**) Schematic of experimental design (see methods). *morcOE* worms were first seeded off hygromycin (P0). P0 worms that lost the extrachromosomal array, identified by loss of muscle mCherry, were moved to either no selection or neomycin selection, then propagated twice to the F2 generation. While F1 worms generally expressed the transgene, silencing was common in F2 worms not maintained on neomycin. Therefore, F2 worms not grown on neomycin were manually separated into transgene-expressing and transgene-silenced populations based on expression of neuronal mCherry. *morcOE* worms maintained on neomycin always retained mCherry expression, but were sick and had low fertility, so few worms remained on neomycin by F2. Hyg = hygromycin, Neo = neomycin, none = no selection. (**E**) Expression changes by RNA-seq of CSR-1 targets vs. non-targets in the three populations of *morcOE* F2 worms assayed: neomycin, no selection + mCherry, no selection – mCherry, see (D). (**F**) Expression changes by RNA-seq of genes binned by wild-type MORC-1 occupancy at TSS (Fig. 2D), in the three populations of *morcOE* F2 worms assayed. (E-F) Expression change shown is the log_2_(fold change) value estimated by DESeq2 (*35*). Effect size measured using Cohen’s d. n.e. = no/minimal effect (|d| < 0.2), * = |d| > 0.2, ** = |d| > 0.5, *** = |d| > 0.9, **** = |d| > 1.5. Note that all these comparisons give p ∼ 0 by Student’s t-test, due to large sample size.

To examine the effect of *morc-1* overexpression on animal fertility, we measured the fertility of individual worms from three populations: non-transgenic animals, *morcOE* worms overexpressing *morc-1*, and *morcOE* worms that have silenced *morc-1* (as assessed by phenotypic indicators of silencing, see Fig. 3B). Although all *morcOE* worms showed reduced fertility compared to wild-type, those that had silenced the *morc-1* transgene were significantly more fertile than worms still expressing the *morc-1* transgene (Fig. 3C). As these two *morcOE* populations are genetically identical and differ only by the expression of the integrated *morc-1* transgene, these data indicate that MORC-1 overexpression alone is sufficient to substantially reduce fertility.

We also examined the effect of MORC-1 overexpression on gene expression by performing RNA-seq on *morcOE* worms at the P0, F1, and F2 generations (Fig. 3D). One population of *morcOE* worms was transferred to neomycin to select against silencing of the integrated *morc-1* transgene, while a second population was maintained on non-selective media. *morcOE* worms on non-selective media began to show loss of transgene expression by the F2 generation (Fig. 3D). We manually separated F2 worms expressing the *morc-1* transgene from those that had silenced it, using the loss of neuronal mCherry expression as a marker for silencing (Fig. 3D). Whereas endogenous *morc-1* expression was similar in all samples, expression of the *morc-1* transgene was only detectable in *morcOE* worms positive for the mCherry signal, but not in F2 worms that had silenced mCherry (fig. S16A-C, see supplemental text), as expected. Consistent with our model, genes downregulated in *morc-1* transgene-expressing samples were highly enriched for germline-expressed genes (fig. S17A-C). In contrast, *morc-1* transgene-silenced samples had few downregulated genes, which were not enriched for any specific tissue or gene ontology terms (fig. S17A-C). Similarly, CSR-1 targets and other genes highly bound by MORC-1 in the wild-type germline were downregulated in *morc-1* transgene-expressing samples, but this effect was substantially reduced in the transgene-silenced samples (Fig. 3E-F, fig. S18A-C). Taken together, these data indicate that MORC-1 overexpression in the germline is sufficient to downregulate the expression of germline-expressed genes.

## Discussion

The prevailing model of CSR-1-mediated gene licensing proposes that CSR-1 directly promotes gene expression, albeit largely through an uncharacterized mechanism (*3*, *4*). Our results position MORC-1 at the core of the CSR-1-mediated gene licensing mechanism, where MORC-1 acts downstream of CSR-1 to tune levels of germline expressed genes. Because CSR-1 robustly slices *morc-1* mRNA, MORC-1 is overexpressed 3–5-fold in *csr-1* mutants (Fig. 1 and (*10*)), where it accumulates at its target genes in excess and causes downregulation. Our data indicate that MORC-1 binds highly expressed genes, which, in the germline, coincides with CSR-1 target genes (*3*). Thus, the primary outcome of MORC-1 overexpression in the germline is downregulation of CSR-1 target genes, suggesting that the mechanism behind CSR-1 germline gene licensing mainly depends on CSR-1-mediated repression of *morc-1*. Because CSR-1 targets represent nearly all germline expressed genes (*3*), we postulate that the sterility associated with *csr-1* mutants is largely a result of global downregulation of germline gene expression by MORC-1. This hypothesis is supported by our finding that *morc-1* mutants can partially rescue *csr-1* mutant sterility and that MORC-1 overexpression alone also induces sterility and downregulation of germline gene expression, including CSR-1 licensed targets.

If CSR-1 does not directly function to promote the expression of germline transcripts, why then are CSR-1-bound 22G-RNAs complementary to the set of germline expressed genes? Interestingly, a recent report demonstrates that CSR-1 is required to silence maternally deposited transcripts in the somatic blastomeres of early embryos (*27*). Since all somatic blastomeres contain transcripts that originate in the maternal germline, it is necessary to clear them to allow for proper zygotic genome activation and somatic development. CSR-1 performs this critical function in embryos by slicing maternally-derived transcripts (*27*). This raises the intriguing possibility that in the germline prior to fertilization, CSR-1 associates with 22G-RNAs complementary to germline targets for the purpose of marking them as ‘germline-lineage’, allowing CSR-1 to later use that ‘memory’ to cleave these targets post-fertilization. More research is needed to understand the function of these CSR-1-bound 22G-RNAs, as well as how the 22G-RNAs generated from germline-expressed transcripts are properly sorted and loaded into CSR-1.

Our results also shed further light on the mechanism of action of MORC-1. By studying MORC-1 localization by ChIP-seq, we found that MORC-1 is strongly enriched at gene promoters, with the most highly expressed genes showing the highest localization of MORC-1. *In vitro*, MORC-1 binds in a non-sequence-specific manner, prefers naked DNA, and is inhibited by competing nucleosomes (*16*). This fits well with the MORC-1 localization pattern described here, where we see strong enrichment of MORC-1 in regions of more accessible chromatin, like the promoters of highly expressed genes. Interestingly, similar promoter binding patterns have been observed for plant and mammalian MORCs, although the function of MORCs at these sites is still not clear (*28–30*). When overexpressed, MORC-1 accumulates further at its targets in a manner proportional to the initial amount of MORC-1. This is consistent with *in vitro* data showing that MORC-1, once bound to DNA, can stimulate binding of other MORC-1 molecules to adjacent stretches of DNA (*16*). The over-accumulation of MORC-1 over gene promoters and bodies leads to gene downregulation, possibly by recruiting H3K9me3 methyltransferases as in human MORC2 (*14*), given the strong increase in H3K9me3 at these sites (Fig. 2F), or possibly through direct chromatin compaction (*16*). Overexpression of MORCs is a hallmark of several cancers (*29*, *31–34*), and missense mutations in MORC2 can cause developmental disorders in humans by hyperactivating the HUSH complex (*14*). Thus, correct MORC dosage appears to be critical for normal gene expression and development across a variety of systems. Further work to understand the consequences of MORC overexpression in other systems, and particularly at gene promoters where the function of MORC remains unclear, should be informative.

In summary, our results demonstrate that MORC-1 is a key component of the CSR-1 gene licensing mechanism, acting to downregulate germline genes in *csr-1* mutants. The viability of germ cells depends on proper MORC-1 expression, which is regulated by CSR-1-mediated slicing of *morc-1* transcripts.

## Acknowledgments

We thank Mindy Clark and Rebecca Tay for extensive discussions and help throughout the entirety of this project. We would like to acknowledge all members of the Kim and Jacobsen Labs for additional insights and discussions. Some strains were provided by the *Caenorhabditis* Genetics Center, which is funded by the NIH Office of Research Infrastructure Programs (P40 OD010440).

## Funding

National Institutes of Health grant R01 HD109667-01 (JKK)

National Institutes of Health grant F32GM136115 (CLP)

W. M. Keck Foundation (SEJ)

SEJ is an Investigator of the Howard Hughes Medical Institute

## Author contributions

Conceptualization: JAK, CLP, NEW, SEJ, JKK

Methodology: JAK, CLP, NEW, NM, DF, VM, SEM, CFJ, SEJ, JKK

Investigation: JAK, CLP, NEW, NM, SF, VM, AV, AFA, CX, KI, SEM

Visualization: JAK, CLP

Funding acquisition: SEJ, JKK

Project administration: SEJ, JKK

Supervision: CFJ, SEJ, JKK

Writing – original draft: JAK, CLP

Writing – review & editing: JAK, CLP, SEJ, JKK

## Data and materials availability

All raw sequencing data generated for this work have been deposited to the GEO database with accession number GSE254933. Data used for analyses, including fertility assays and sequencing data, are all available in the supplementary materials.

## Supplementary Materials

### Materials and Methods

#### Experimental model and subject details

*C. elegans* strains were maintained using standard procedures (*36*) at 15°C or 20°C unless indicated otherwise. In all cases, Bristol N2 strain was used as the wild-type control. Worms were fed OP50 *E. coli* for all experiments except ChIP-seq of purified germline nuclei and those involving RNAi. Worms for germ cell ChIP were fed HB101 *E. coli* and worms for RNAi experiments were fed with HT115 *E. coli*.

#### Construction of transgenic animals

All CRISPR strains were generated according to standard procedure, as described (*37*). The MORC-1 overexpression strain (*morcOE*) was created by first generating an extrachromosomal array by injecting into the *unc-119(ed3)* strain: 1) a piRNA that silences the recoded, codon-optimized *morc-1::3xflag* transgene (*morc-1(co)::3xflag*) in the germline via the piRNA pathway (*26*), 2) reagents to generate targeted array integrations (Cas9 and sgRNA), 3) a Hygro^R^ transgene for selection, and 4) *Pmlc-1::mCherry*. These initial transgenic worms were then injected with 1) the recoded *morc-1* transgene containing PATCs (*Pmex-5::morc-1(PATC)::3xFlag::gpd-2::ce-gfp*), 2) 1kb ladder DNA, 3) *Prab-3::mCherry*, 4) Neomycin^R^, and 5) a fragment for targeted array integration into the *unc-119(ed)* locus using the MosTi single-copy integration method (*25*). The full genotype of *morcOE* is: kstSi107pSEM417 *(Pmex-5::morc-1(PATC)::3xFlag::gpd-2::ce-gfp*), pSEM371 (*unc-119* integration fragment), pGH8 (*Prab-3::mCherry*), pCFJ594 (NeoR)] III; kstEx75[pCFJ2474 (*Psmu-2::Cas9(PATC)::gpd-2::tagRFP(myr),* pSEM376 (sgRNA *ce-unc-119* locus), T1636 (*morc-1* recoded piRNAi), pSEM235 *(Pmlc-1::mCherry*), pCFJ782 (Hygro), 1 kb ladder].

#### RNAi assays

Bacterial clones containing the RNAi of interest were grown from the Ahringer RNAi library (*38*) and administered by feeding, as reported previously (*39*). Cultures were inoculated from a single colony, grown for 12–16 hr in Luria Broth with carbenicillin (50 µg/mL) and plated on IPTG-containing plates, then induced for expression overnight at 25°C. Worms were plated after this induction period and grown at 20°C, unless indicated otherwise. In all cases, the empty vector L4440 was used as a negative control.

#### Auxin assays

In experiments in which auxin (3-indoleacetic acid) (Sigma-Aldrich) was used, NGM plates were poured containing the indicated concentration of auxin. In ChIP-seq experiments in which worms were grown in liquid culture, the auxin was added directly to the cultures.

#### Western blotting

For western blots, worms were lysed in Tris-glycine SDS sample buffer (ThermoFisher), run on an 8-16% Novex WedgeWell Tris-Glycine pre-cast gel (ThermoFisher), then transferred to a PVDF membrane (Millipore) using a Trans-Blot Turbo Transfer System (Bio-Rad). Primary antibodies used were Sigma-Aldrich F1804 (anti-FLAG) at 1:1,000, Abcam ab1791 (anti-H3) at 1:15,000, and anti-CSR-1 (*16*) at 1:2,000. Secondary antibodies used for western blots developed in a BioRad ChemiDoc Touch system and exposed using Pierce ECL (ThermoFisher) were GE Healthcare NA931 (sheep anti-mouse) at 1:2,000 and Jackson Laboratories 111035045 (goat anti-rabbit) at 1:15,000 when used with anti-H3 and 1:10,000 when used with anti-CSR-1 antibodies. Western blots developed using a LI-COR Odyssey Fc were done according to the manufacturer’s instructions using Odyssey Blocking Buffer and IRDye secondary antibodies at 1:15,000 (LI-COR).

#### Single generation and transgenerational fertility assays

Gravid worms were hypochlorite treated and their progeny, the P0 generation, were shifted to the appropriate temperature (25°C was used if not specifically stated) for all assays. Either the P0 generation or their progeny, the F1 generation, where indicated, were singled at the L2–L3 stage and their total progeny were counted. For transgenerational fertility assays, in addition to singling at the L2–L3 stage, ∼20 F1 worms were transferred to a single additional propagation plate so that their progeny could be singled at the subsequent generation.

#### DIC imaging of worms

DIC images of worms were acquired on the Zeiss Axio Zoom V16 Fluorescence Stereo Microscope.

#### Him assays

Worms were synchronized via hypochlorite treatment and their progeny were plated at 20°C. At the L4 stage, 10 hermaphrodites were transferred to a new plate. The sex of their progeny was then scored.

#### Immunofluorescence of the *C. elegans* germline

Gravid adult *C. elegans* were dissected in egg buffer (118 mM NaCl, 48 mM KCl, 2 mM EDTA, 0.5 mM EGTA, 25 mM HEPES [pH 7.4]), containing 15 mM sodium azide and 0.1% Tween-20 then fixed in 1% formaldehyde in egg buffer for 10 sec followed by a 1 min methanol fixation at -20°C. Primary mouse anti-FLAG antibody (Sigma F1804) was used at 1:100 and rabbit anti-CSR-1 (*16*) was used at 1:200 in normal goat serum and PBST. The secondary antibody was used at 1:300 (Invitrogen AlexaFluor 555 goat anti-mouse and 488 goat anti-rabbit) in PBST. All washes and staining were performed in suspension. Germlines were stained with DAPI (0.5 mg/mL) then mounted with Vectashield (Vectorlabs H-1000). Images were acquired on a Zeiss LSM700 confocal microscope at 63x magnification. Image processing was performed using Zen SP5 software.

#### Chromatin immunoprecipitation on purified germline nuclei

A synchronized population of worms was obtained by hypochlorite treatment of gravid adult worms followed by overnight nutation of the embryos in M9. Worms were grown in liquid culture at 20°C according to the protocol described in (*40*) and collected at 56 hours (young adult stage). *morc-1::3xflag; aid::csr-1; psun-1::TIR1* (*aid::csr-*1) worms were grown in the presence of 50 µM auxin beginning at the L1 stage. At the 56 hour time point, the worms were cleansed via sucrose floatation (*40*). They were then live-crosslinked in 2.6% formaldehyde for 30 min at room temperature with nutation. The crosslinker was quenched with a 5 min nutation in glycine at a final concentration of 125 mM. Worms were washed in water and flash frozen in liquid nitrogen as ∼1 mL pellets. Germline nuclei were purified according to Han, *et al*., 2019 (*22*) with some modifications. In brief, frozen worms were ground in an MM400 Mixer Mill homogenizer (Retsch) for 2 rounds of 15 sec at a frequency of 30^-1^ sec. Frozen worm powder from each pellet was resuspended in 10 mL of nuclear purification buffer (50 mM HEPES (pH 7.5), 40 mM NaCl, 90 mM KCl, 2 mM EDTA, 0.5 mM EGTA, 0.2 mM DTT, 0.5 mM PMSF, 0.5 mM spermidine, 0.25 mM spermine, and 1 cOmplete ULTRA tablet (Roche) per 25 mL buffer) and allowed to chill on ice for 5 min. Resuspension was aided by 30 sec of vortex at max speed. One additional round of 5 min on ice followed by 30 sec max speed vortexing was performed. Samples were then spun at 30 x g for 5 min at 4°C. The supernatant was successively passed through two 40 µm filters (pluriSelect), then two 30 µm filters (pluriSelect), and finally two 20 µm filters (pluriSelect). Nuclei were pelleted at 2,400 x g for 6 min at 4°C and the supernatant removed. Purified germline nuclei were resuspended in 1 mL nuclear purification buffer, transferred to LoBind tubes (Eppendorf), re-pelleted at 1,500 x g for 5 min at 4°C and flash frozen after removing the supernatant. Nuclei were resuspended in 1x RIPA buffer (1x PBS, 1% NP-40, 0.5% sodium deoxycholate, 0.1% SDS) and nutated for 10 min at 4°C. Chromatin was sheared to a length of 100–500 bp using a Bioruptor Pico water bath sonicator (Diagenode) for three 3-min cycles, 30 sec on/off. Crosslinked chromatin was immunoprecipitated overnight at 4°C with 2 µg Flag antibody (Sigma-Aldrich, F1804) and then for 2 hr with 50 µL Protein G Dynabeads (Invitrogen). Immunoprecipitated material was washed 3x in LiCl buffer (100 mM Tris-Cl pH 7.5, 500 mM LiCl, 1% NP-40, 1% sodium deoxycholate). The crosslinking was then reversed in worm lysis buffer (0.1 M Tris-Cl pH 7.5, 0.1 M NaCl, 50 mM EDTA, 1% SDS) with 6.8 µM Proteinase K for at least 4 hr at 65°C in a Thermomixer (Eppendorf). DNA was extracted by phenol-chloroform and dissolved in TE buffer. RNAse A (Invitrogen) treatment was performed for at least 2 hr at 37°C. For each immunoprecipitated sample, an input library was generated from 10% of the chromatin pre-IP and was de-crosslinked and extracted in parallel to the immunoprecipitated samples. Precipitated DNA was then used for library preparation using the Ovation Ultra Low System V2 kit (NuGEN) according to the manufacturer’s instructions and then sequenced on a NovaSeq 6000 Sequencer (Illumina).

#### Chromatin immunoprecipitation on whole worms

Worms previously maintained at 20°C were hypochlorite treated and shifted to 25°C; these worms became the “P0” population. Worms were propagated at 25°C for four successive generations (F1–F4); each generation was synchronized by hypochlorite treatment. When *csr-1* RNAi was used, it was fed to the worms beginning at the L1 stage only for the single generation prior to worm sample collection. Chromatin sonication, immunoprecipitation, de-crosslinking, and DNA extraction were performed as previously described (*15*). Antibodies used were Abcam 8898 (H3K9me3) and WAKO 300-95289 (H3K36me3). Precipitated DNA was then used for library preparation using the Ovation Ultra Low System V2 kit (NuGEN) according to the manufacturer’s instructions and then sequenced on a NovaSeq 6000 Sequencer (Illumina).

#### RNA extraction

Worms were collected in TriReagent (ThermoFisher) and subjected to three freeze-thaw cycles. Phase separation was achieved via 1-bromo-3-chloropropane (BCP) and the aqueous phase was precipitated in isopropanol at -80°C for 2 hrs. To pellet RNA, samples were centrifuged at 21,000 x g for 30 min at 4°C. The pellet was washed three times in 75% ethanol then resuspended in water.

#### Quantitative RT-PCR

cDNA for quantitation of mRNA levels was made from 500 ng of total RNA using Multiscribe Reverse Transcriptase (Applied Biosystems) and random hexamer primers in an Eppendorf Mastercycler Pro S6325 (Eppendorf). qPCR for mRNA levels was performed with Absolute Blue SYBR Green (ThermoFisher) and normalized to *eft-2* or *him-3* (in experiments using the MORC-1 overexpression strain (*morcOE*)) using a CFX63 Real Time System Thermocycler (Bio-Rad). Specific primers used to measure mRNA levels are as follows: *morc-1:* GAAGCTGTGTCAAATGTGCCG and GAGAGTCGGACGATGATGGTG; codon-optimized *morc-1* transgene in *morcOE*: GAAGCTTGAGAAGGCCTCTGT and CGAGCCATTCCAAGACCATCA; *eft-2:* ACGCTCGTGATGAGTTCAAG and ATTTGGTCCAGTTCCGTCTG; *him-3:* CGACGGATTGAGAGATGCGA and CGTTCGTGTCGATTCCGTTAT.

#### MORC-1 overexpression experimental setup

*MorcOE* worms were maintained continuously on 4 mg/mL hygromycin to select for worms inheriting the extrachromosomal array and 25 mg/mL neomycin (G418 Goldbio). To initiate an assay, *morcOE* worms were hypochlorite treated and their progeny, the P0 generation, were seeded on NGM plates without antibiotic. P0 worms that lost the extrachromosomal array, by visual inspection for the absence of muscle-expressed mCherry (from *Pmlc-1::mCherry*), were singled. Loss of the extrachromosomal array was confirmed by genotyping (using primers aattttccagTCCAAGGCCG and GTCTGGGTTCCCTCGTATGG). F1 progeny were then singled onto NGM plates that either contained 25 mg/mL neomycin or no antibiotic. In addition to the singled worms, a plate of >30 worms of each condition (+/-neomycin) were plated for RNA extraction. The phenotype of the singled hermaphrodite was scored for uncoordinated (Unc) movement or wild-type movement and for neuronal mCherry-ON or mCherry-OFF by visual inspection under a fluorescence dissecting microscope (Leica). Fertility of each singled worm was measured. When the progeny of the worms plated for RNA extraction reached the adult stage, they were collected and stored in TriReagent (ThermoFisher). This process was repeated at the F2 generation. However, at the F2 generation, worms that had been reared without antibiotic were showing significant silencing of the neuronal mCherry (*Prab-3::mCherry*); therefore, for the RNA collection samples, the worms were first manually separated according to their expression state of the neuronal mCherry (ON vs OFF).

#### RNA-sequencing library preparation and sequencing

RNA-seq libraries were generated using the KAPA stranded mRNA-seq kit (Roche) according to the manufacturer’s instructions. Small RNA-seq libraries were made using the NEBNext Multiplex Small RNA Library Prep Set (NEB) according to the manufacturer’s instructions. All RNA-seq libraries were sequenced on a NovaSeq 6000 Sequencer (Illumina).

#### ATAC-seq

ATAC-seq was performed on adult wild-type (N2) worms grown at 25°C. ATAC-seq was performed as previously described (*41*) and sequenced on a NovaSeq 6000 Sequencer (Illumina).

#### Sequencing data analysis

For published datasets reanalyzed for this study (Table S2), raw data were re-downloaded from the GEO database and processed using the same pipeline described here. All sequencing data were first checked for quality using fastqc v0.11.8 (*42*). Reads were filtered and trimmed to remove poor quality sequences and adapter sequences using Trim Galore v0.6.7 (*43*) with options --stringency 3 -q 25 --length 20. Reads were aligned to ce10/WBcel215 with the WS230 annotations, obtained from WormBase (https://wormbase.org). RNA-seq reads were aligned using STAR v2.7.9a (*44*) with options --outFilterMismatchNoverReadLmax 0.05 -- alignIntronMin 70 --alignIntronMax 5000 --alignMatesGapMax 100000 --outFilterIntronMotifs RemoveNoncanonical --alignEndsType EndToEnd, while ChIP-seq and ATAC-seq reads were aligned using bowtie2 v2.3.4.3 (*45*) with options -N 0 -L 22 (-X 500 for paired-end data). PCR duplicates were removed using MarkDuplicates from the Picard tools suite (*46*). Alignment statistics for all libraries generated for this study are available in Data S1.

#### ATAC-seq data analysis

Aligned reads were analyzed using Genrich v.0.5 (*47*) to identify cut sites, excluding the mitochondrial chromosome. Cut site pileups from Genrich were used to make metaplots over gene promoters using DeepTools (*48*) computeMatrix and plotHeatmap functions.

#### ChIP-seq data analysis

Fragment size was estimated using the run_spp function from the phantompeakqualtools (*49*, *50*) suite. Fragment size estimates averaged to approximately 175 bp across all samples, and this estimate was used for all analyses. Read coverage tracks for each sample were generated using the bamCoverage function from DeepTools (*48*) suite v.3.5.1, using options --extendReads 175 - -binSize 10 and --normalizeUsing CPM, and excluding the mitochondrial chromosome. Additionally, coverage tracks of log_2_(IP/input) were also generated using the DeepTools bamCompare function again with --extendReads 175 --binSize 10 and --normalizeUsing CPM. For conditions with multiple replicates, average log_2_(IP/input) signal tracks were generated by (1) summing all IP tracks together using the DeepTools bigWigMerge function, (2) summing all input tracks together (each IP track has a single matched input track), and (3) calculating log_2_(sum IP / sum input) using DeepTools bigwigCompare. Average signal across specific genomic features (e.g. over promoters, or 1kb bins tiled genome-wide) were obtained using the DeepTools multiBigwigSummary function. The transcriptional start site (TSS) region for a gene was defined as 1,000 bp upstream to 400 bp downstream.

Signal peaks were identified using the MACS2(*51*) v.2.2.7.1 callpeak function, with parameters - g 93260000 --broad -f BAM --nomodel --extsize 175. Peaks were called between each replicate IP and its matched input. For conditions with multiple replicates, consensus peaks were identified by first merging peaks across all replicates using bedtools (*52*) merge (‘union peaks’). The union set of peaks was then filtered to only keep those that overlap at least 50% with each individual replicate: bedtools intersect -wa -a union_peaks.bed -b rep1_peaks.bed -f 0.5 -F 0.5 -e, producing the final set of consensus peaks. To calculate peak overlap with various genomic features (PCGs, promoters, repeats, TEs, etc.), BED files containing these different regions were first combined into a single file, and an extra column was added containing an integer ranking, such that TEs > PCG exons > PCG introns > non-PCG bodies > repeats > PCG promoter (2 kb up) > non-PCG promoter (2 kb upstream). Intervals in the consensus peak files were intersected with the regions BED file using bedtools intersect with options -a consensus_peaks.bed -b regions.bed -wao -f 0.1. Peaks overlapping multiple regions in the BED file were ranked by the ranking column, so that if, for example, both a TE and a PCG exon overlapped, the peak was assigned to ‘TE’. All peaks not overlapping any regions were assigned to category ‘other’.

Metaplots and heatmaps of ChIP-seq signal over genomic features/intervals were obtained using the DeepTools computeMatrix function, followed by plotProfile and/or plotHeatmap. K-means clustering of heatmap rows was performed using the --kmeans option in plotHeatmap, with optimal *k* chosen by visual inspection. To bin genes based on MORC-1 signal at the TSS in wild-type, the difference in average log2(IP/input) signal between wild-type MORC-1 and the no-FLAG control over the TSS was calculated. The bins “none”, “weak”, “medium”, “strong” were assigned to genes with signal difference <= 0, (0,0.5], (0.5,1], and > 1 respectively.

#### RNA-seq data analysis

RNA-seq coverage tracks were generated using DeepTools (*48*) bamCoverage with options -- binSize 10 --normalizeUsing CPM. Counts over genes were obtained using the htseq-count function from the HTSeq python package (*53*) v2.0.2 with options --nonunique none -m intersection-strict, and using the WS230 *C. elegans* annotations. Raw counts from each sample were combined into a counts matrix, and DESeq2 (*35*) was used to estimate expression changes between conditions (log_2_(fold change)), and to identify significantly differentially expressed genes. For *morcOE* samples, each condition was compared to the non-transgenic control.

Metaplots and heatmaps of ChIP-seq signal over genomic features/intervals were obtained using the DeepTools computeMatrix function, followed by plotProfile and/or plotHeatmap. TPM estimates were obtained using StringTie v.2.1.6 (*54*).

#### sRNA-seq data analysis

Reads were first trimmed to remove adapter sequences using Trim Galore (*43*) with options -- illumina --max_length 35 --length 18 -q 0 --stringency 3, retaining only reads between 18-35 bp post-trimming. Reads were further split according to size: 22 nt (CSR-1 and WAGO siRNAs plus miRNAs), 21 nt (PRG-1 piRNAs) and 26 nt (ERGO-1 and ALG-3/4-dependent siRNAs), and aligned to ce10 using bowtie2 (*45*) in --end-to-end --very-sensitive mode. Uniquely mapped reads were extracted (mapQ >= 2), and converted into strand-specific coverage tracks using DeepTools (*48*) bamCoverage --binSize 10 --normalizeUsing CPM --filterRNAstrand [forward/reverse]. Counts over genes were obtained using the htseq-count function from the HTSeq python package (*53*) v2.0.2 with options --nonunique none -m intersection-strict, and using the WS230 *C. elegans* annotations, and log_2_(fold change) estimates between wild-type and mutants were estimated using DESeq2 (*35*).

#### Gene Ontology Analysis

Gene ontology analysis for enriched GO-terms was performed using DAVID (*55*) (fig. S19B). Tissue enrichment analysis (fig. S10C, S17B) was performed using the WormBase Enrichment Analysis tool (*56*).

#### Genome Browser Images

Images of signal tracks were taken using the Integrative Genomics Viewer (IGV) (*57*).

#### Published Datasets

Additional published datasets used in this study are shown in Table S2. Some of these were re-downloaded and reanalyzed using the same analysis pipeline described above, and mapping statistics for these libraries are available in Data S1.

#### Data Plotting

Heatmaps and metaplots were made using DeepTools (*48*). Most other plots were made using Stata version 14 (*58*).

### Supplementary Text

#### *morc-1* does not rescue the *csr-1* Him phenotype

Partial loss of the CSR-1 pathway, which enables worms to produce a small number of progeny but still recapitulates many other *csr-1* defects, also leads to a high incidence of males (Him) phenotype, due to defects in chromosome segregation (*3*). In our RNA-seq data, both *csr-1(G560R)* and *aid::csr-1* showed strong upregulation of a shared set of genes enriched for sperm- and male-related gene ontology terms (fig. S19A-C). We speculated that this could be due to the Him phenotype, and examined the % of male progeny in both of our *csr-1* mutants. Indeed, we observed elevated rates of males in our *csr-1(G560R)*, while *aid::csr-1* could not be evaluated due to low fertility on auxin (fig. S19D), but nonetheless showed the elevated expression of male-related genes by RNA-seq (fig. S19A-C). This suggests that both of our *csr-1* mutants display the Him phenotype.

Unlike the *csr-1* chromatin defects, CSR-1 target gene downregulation, and fertility defects (Fig. 1C, Fig. 2E-F), the Him defect was not rescued in the *csr-1(G560R); morc-1*(*-*) double mutant (fig. S19D). Instead, the defect was mildly enhanced (fig. S19D). Consistent with this, the set of genes that are highly upregulated in *csr-1(G560R)*, which are enriched for male-related genes (fig. S19A-C), were further upregulated in the *csr-1(G560R); morc-1*(*-*) double mutant (fig. S19E). These genes are also mildly upregulated in a published *morc-1* RNA-seq dataset (*15*), suggesting they may normally be repressed by MORC-1 (fig. S19E), which would explain why the *csr-1* Him phenotype is mildly enhanced in the *csr-1(G560R); morc-1*(*-*) double mutant.

These data suggest that the chromosome segregation defects in *csr-1* are in a separate pathway than the defects rescued by *morc-1*(*-*). This is consistent with previous work showing that ectopic upregulation of a different CSR-1 slicing target, *klp-7*, is the primary cause of chromosome segregation defects in *csr-1* (*10*).

#### Additional validation of *morcOE* lines

To validate our RNA-seq data from the *morcOE* lines, we checked the expression of both the integrated transgene and the extrachromosomal array, as well as endogenous *morc-1*, across all our RNA-seq samples (Fig. S16A-C) First, we confirmed that endogenous *morc-1* levels are similar across all of these lines as well as in the non-transgenic control, confirming that the *morcOE* worms have normal expression from the endogenous *morc-1* locus regardless of transgene expression (fig. S16B). We next confirmed that genes from the extrachromosomal array were expressed as expected. The extrachromosomal array contains both the piRNAi construct and the muscle mCherry, and should be lost after the P0 generation, as worms are manually selected for negative muscle mCherry expression (Fig. 3A, see methods). Expression of both genes was only detectable in the P0 generation, as expected (fig. S16C, see methods). Finally, the integrated transgene contains both a neuronal mCherry marker and the codon-optimized *morc-1*. We confirmed that both genes are expressed in all mCherry positive samples, but not in the mCherry negative samples, as expected. More generally, both genes show similar expression patterns, indicating that neuronal mCherry is a good proxy for transgene *morc-1* expression (fig. S16C).

All *morcOE* worms, regardless of the status of *morc-1* transgene expression, shared a set of largely overlapping upregulated genes (fig. S17A-C), indicating that these genes are not upregulated due to *morc-1* overexpression. These genes were not significantly enriched for germline genes or any other significant gene ontology terms (fig. S17B-C).

**Fig. S1.**
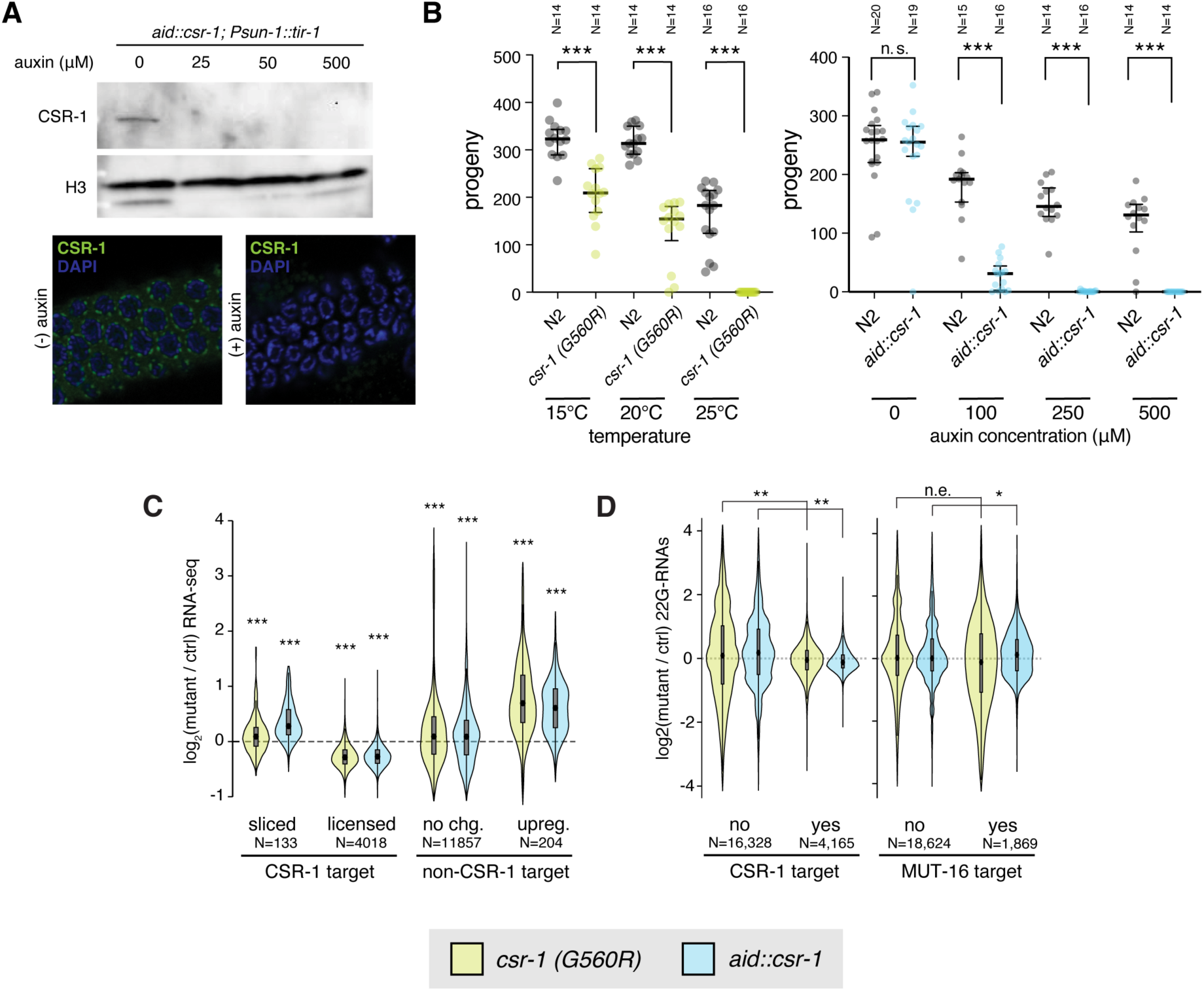
Validation of *csr-1(G560R)* and *aid::csr-1.* (**A**) (top) Western blot for CSR-1 (native antibody) in *aid::csr-1* worms exposed to increasing concentrations of auxin. (bottom) Immunostaining of CSR-1 in dissected germlines from *aid::csr-1* worms on 0 μM [(-) auxin] or 100 μM auxin [(+) auxin]. (**B**) Number of progeny in *csr-1(G560R)* at permissive vs. restrictive temperature (left), and *aid::csr-1* worms exposed to increasing concentrations of auxin vs. wild-type control (right). Each point represents a single worm (n ≥14 worms assayed per genotype/condition). n.s. = p > 0.05, *** = p < 0.001, one-tailed t-test. (**C**) Average RNA-seq expression log_2_ fold change vs. control of *csr-1(G560R)* (control = N2) and *aid::csr-1* (control = *aid::csr-1* without auxin) worms, over CSR-1 sliced targets vs. licensed targets, *csr-1*(*-*) upregulated non-targets (upreg.) vs. all other non-targets (no chg.). Sliced, licensed, unchanged and upregulated genes were identified in Gerson-Gurwitz *et al.* 2016 (*10*). Significance measured by two-tailed t-test, n.s. = not significant, *** = p < 0.0001, with null hypothesis that log_2_(fold change) == 0. (**D**) Average log_2_ fold change of *csr-1(G560R)* and *aid::csr-1* vs. control of 22G-RNAs over CSR-1 targets and MUT-16 targets vs. non-targets (*3*). Effect size measured using Cohen’s d. n.e. = no/minimal effect (|d| < 0.2), * = |d| > 0.2, ** = |d| > 0.5, *** = |d| > 0.9, **** = |d| > 1.5. Note that these comparisons give p ∼ 0 by Student’s t-test due to large sample size.

**Fig. S2.**
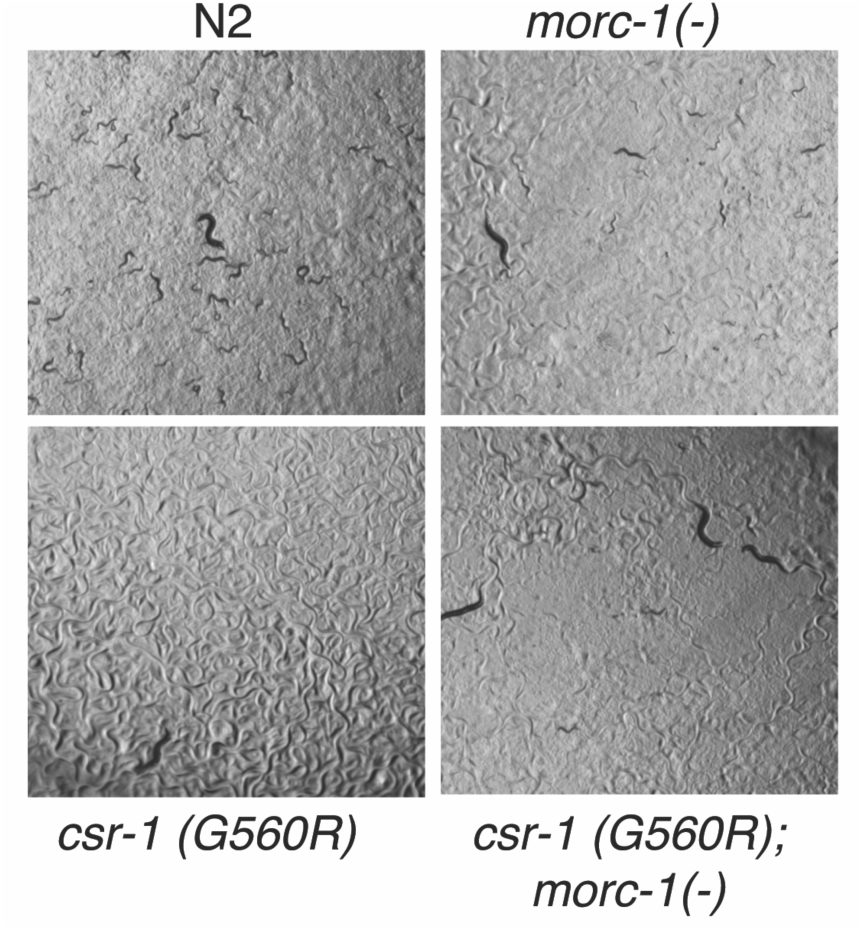
Rescue of *csr-1(G560R)* fertility by *morc-1.* Example DIC images of the parents and progeny of 10 parental worms of each genotype grown at the *csr-1(G560R)* nonpermissive temperature of 25°C.

**Fig. S3.**
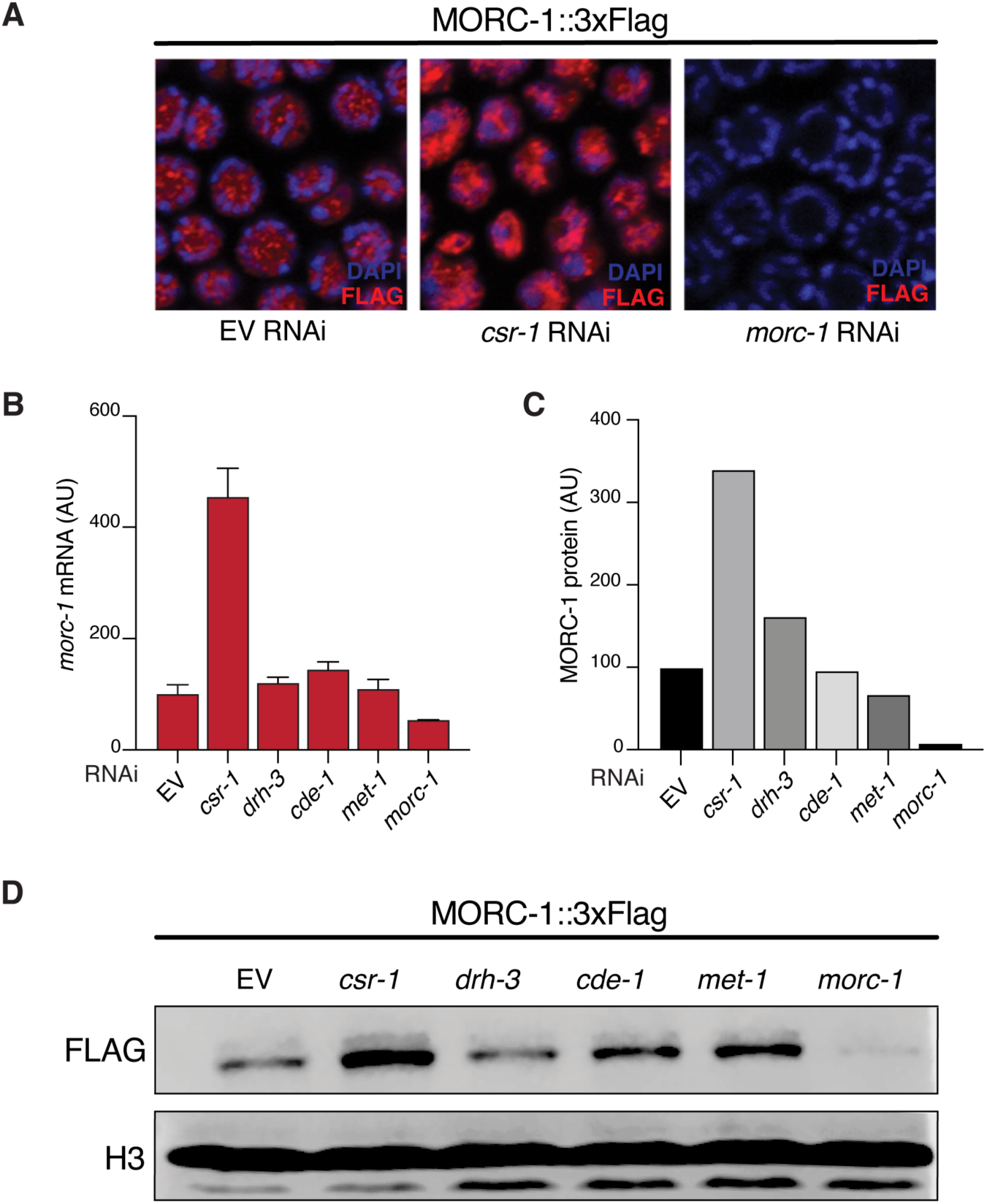
Control of *morc-1* expression by CSR-1 and other members of the CSR-1 sRNA pathway. (**A**) Immunofluorescence of MORC-1::3xFLAG treated with *csr-1* or *morc-1* RNAi compared to empty vector (EV) control, in dissected germlines. FLAG signal shown in red, DAPI in blue. (**B**) *morc-1* mRNA quantification by qPCR for wild-type (N2) worms treated with RNAi against the indicated genes related to the CSR-1 pathway. EV = empty vector RNAi control. Error bars represent standard deviation of two technical replicates. (**C**) Quantification of protein levels in the western blot for MORC-1::3xFLAG shown in (D), in N2 worms treated with RNAi against the indicated genes. (**D**) Western blot corresponding to (C).

**Fig. S4.**
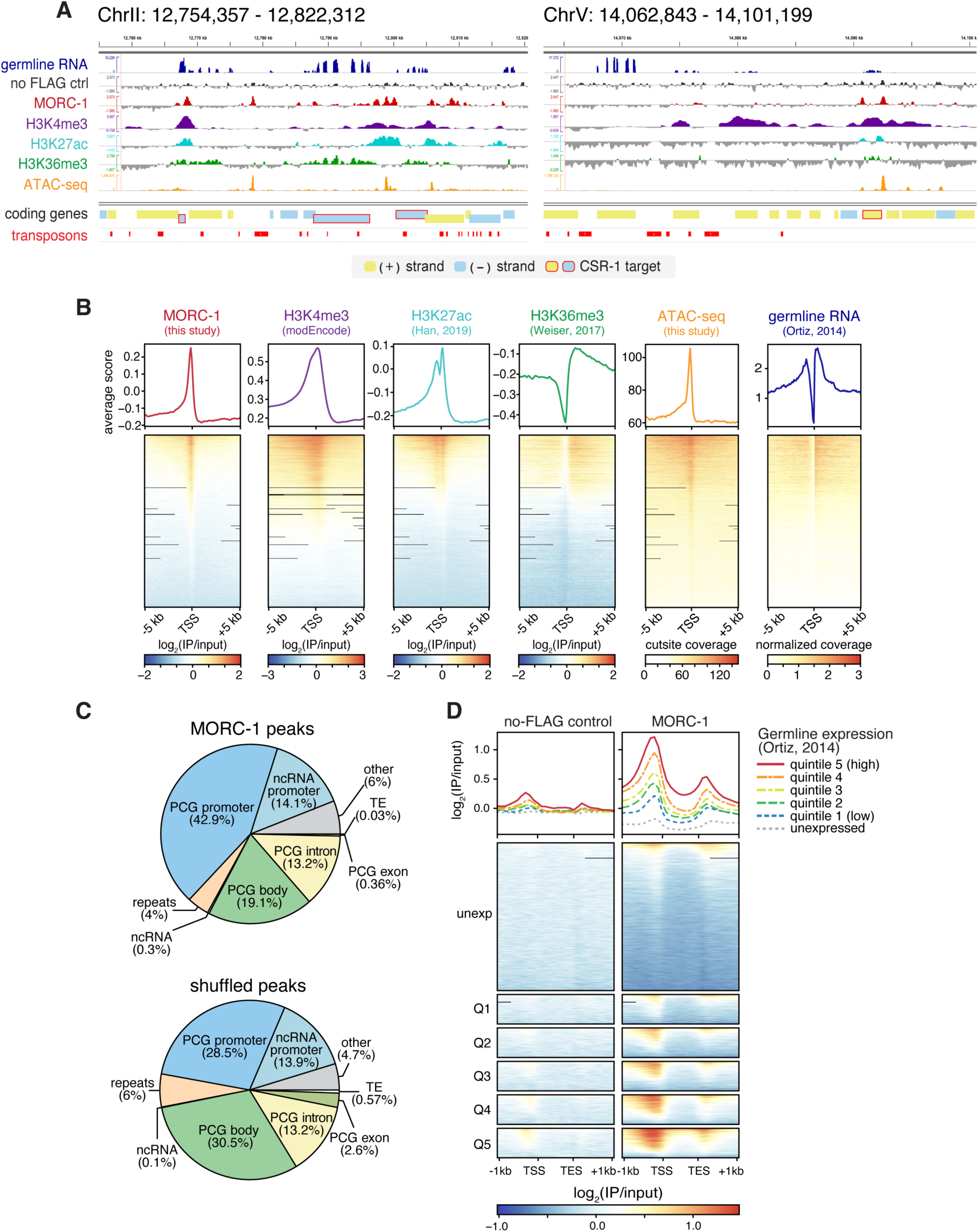
MORC-1 localization in *C. elegans*. (**A**) Example genome browser images showing MORC-1::FLAG localization alongside other published histone modification ChIP-seq datasets (see Table S2), germline expression from Ortiz *et al.* 2014 (*23*), and chromatin accessibility as evaluated by ATAC-seq (this study). (**B**) Metaplots and heatmaps of MORC-1::FLAG, published histone modification ChIP-seq (see Table S2), chromatin accessibility (this study), and germline RNA-seq (*23*) over gene transcriptional start sites (TSSs), ranked by MORC-1::FLAG levels (leftmost heatmap). All heatmaps have same row order. (A-B) For ChIP-seq, the plotted values are log IP signal normalized to input (log_2_(IP/input)), for ATAC-seq the plotted values are the number of ATAC-seq cut sites (see methods), and for RNA-seq the values shown are normalized coverage. (**C**) Distribution of MORC-1 peaks across different genome annotations. PCG = protein-coding gene. Promoters are defined as the region from 2 kb upstream of TSS to the TSS. Peaks in PCG bodies preferentially over introns or exons (> 70% of length of peak) are listed explicitly as ‘PCG intron’ or ‘PCG exon’, while all other peaks in PCG bodies are listed as ‘PCG body.’ True peak distribution was compared to the distribution of randomly shuffled peaks. (**D**) Heatmaps of ChIP-seq signal from no-FLAG control (left) or MORC-1::FLAG (right), over protein-coding genes binned into quintiles by germline expression level from Ortiz *et al.* 2014 (*23*). Genes detected in Ortiz *et al.* as germline-expressed were divided into quintiles Q1-Q5 based on average TPM levels in the Ortiz *et al.* RNA-seq, with Q5 containing the most highly expressed genes, while unexpressed genes (unexp) were kept as a separate group.

**Fig. S5.**
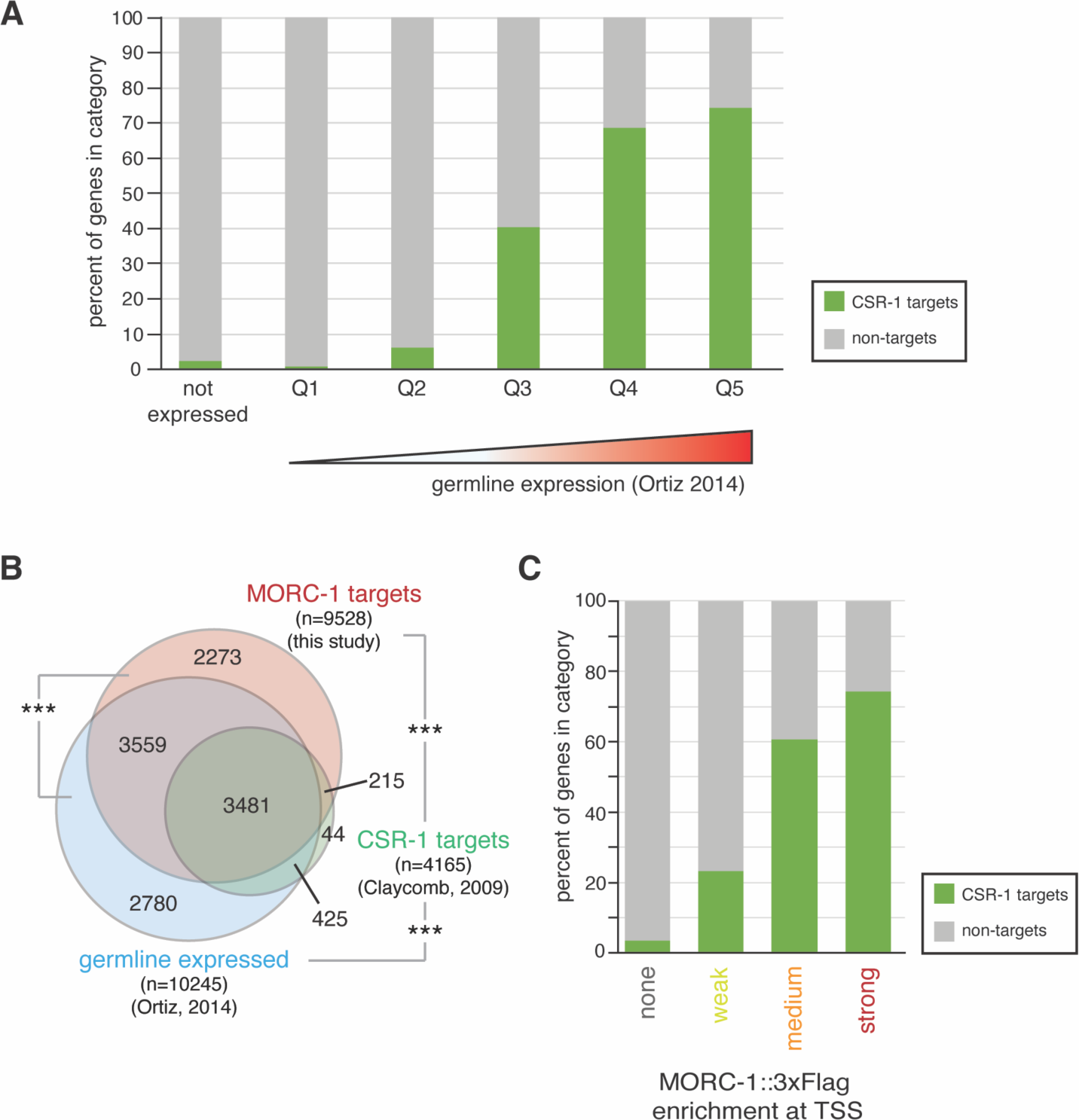
CSR-1 targets are highly expressed in the germline and strongly bound by MORC-1. (**A**) Percent of genes binned by germline expression level that are CSR-1 targets. Genes detected in Ortiz *et al.* 2014 (*23*) as germline-expressed were divided into quintiles based on average TPM levels in the Ortiz *et al.* RNA-seq, with Q5 containing the most highly expressed genes. (**B**) Overlap between CSR-1 targets (green) from Claycomb *et al.* 2009 (*3*), germline expressed genes (blue) from Ortiz *et al.* 2014 (*23*), and MORC-1 targets (red, this study) identified in this study based on MORC-1 enrichment over the promoter and TSS. *** = p ∼ 0, hypergeometric test. (**C**) Percent of genes binned by MORC-1 enrichment at the TSS (same bins as Fig. 2D) that are CSR-1 targets identified in Claycomb *et al.* 2009.

**Fig. S6.**
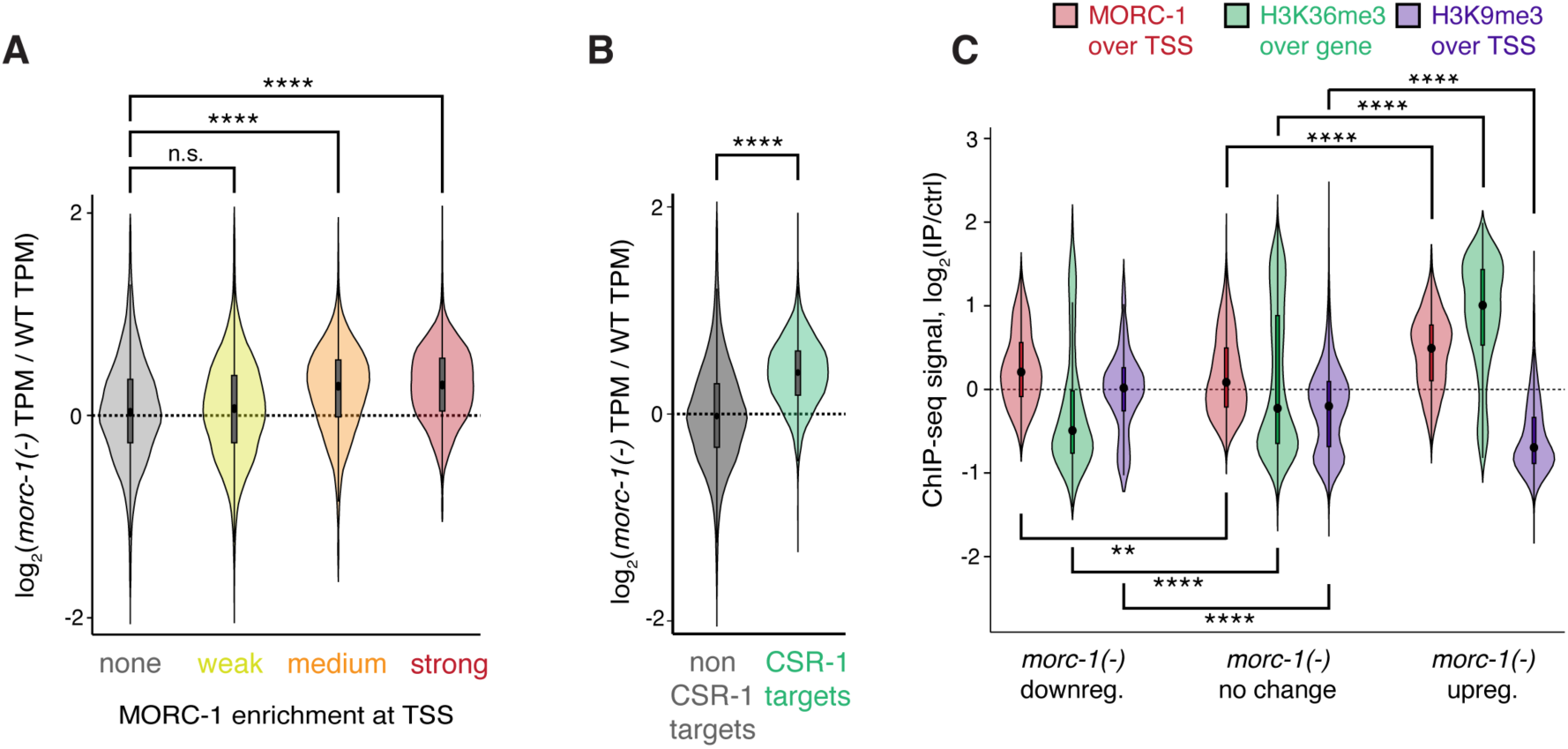
Relationship between MORC-1 occupancy and *morc-1*(*-*) expression changes. (**A-B**) Expression change in *morc-1*(*-*) from Weiser *et al.* 2017 (*15*), as a function of MORC-1 levels at the transcription start site (TSS) in wild-type animals (A), or CSR-1 targets vs. non-targets (B). A small number of genes (N=68) with y-values outside of [-2,2] not shown. (**C**) Wild-type ChIP-seq signal (log_2_(IP/input)) of MORC-1::3xFLAG (this study) over TSS, H3K36me3 (Weiser *et al.*) over gene body, and H3K9me3 (Weiser *et al.*) over TSS, comparing genes significantly upregulated (N=1,302), downregulated (N=403), or unchanged in *morc-1*(*-*) based on Weiser *et al.* 2017 RNA-seq. All data from Weiser *et al.* are from the F4 generation of *morc-1*(*-*) worms maintained at 25°C. (A-C) Significance by two-tailed t-test: n.s. = not significant, * = p < 0.01, ** = p < 0.001, *** = p < 0.0001, **** = p < 0.00001.

**Fig. S7.**
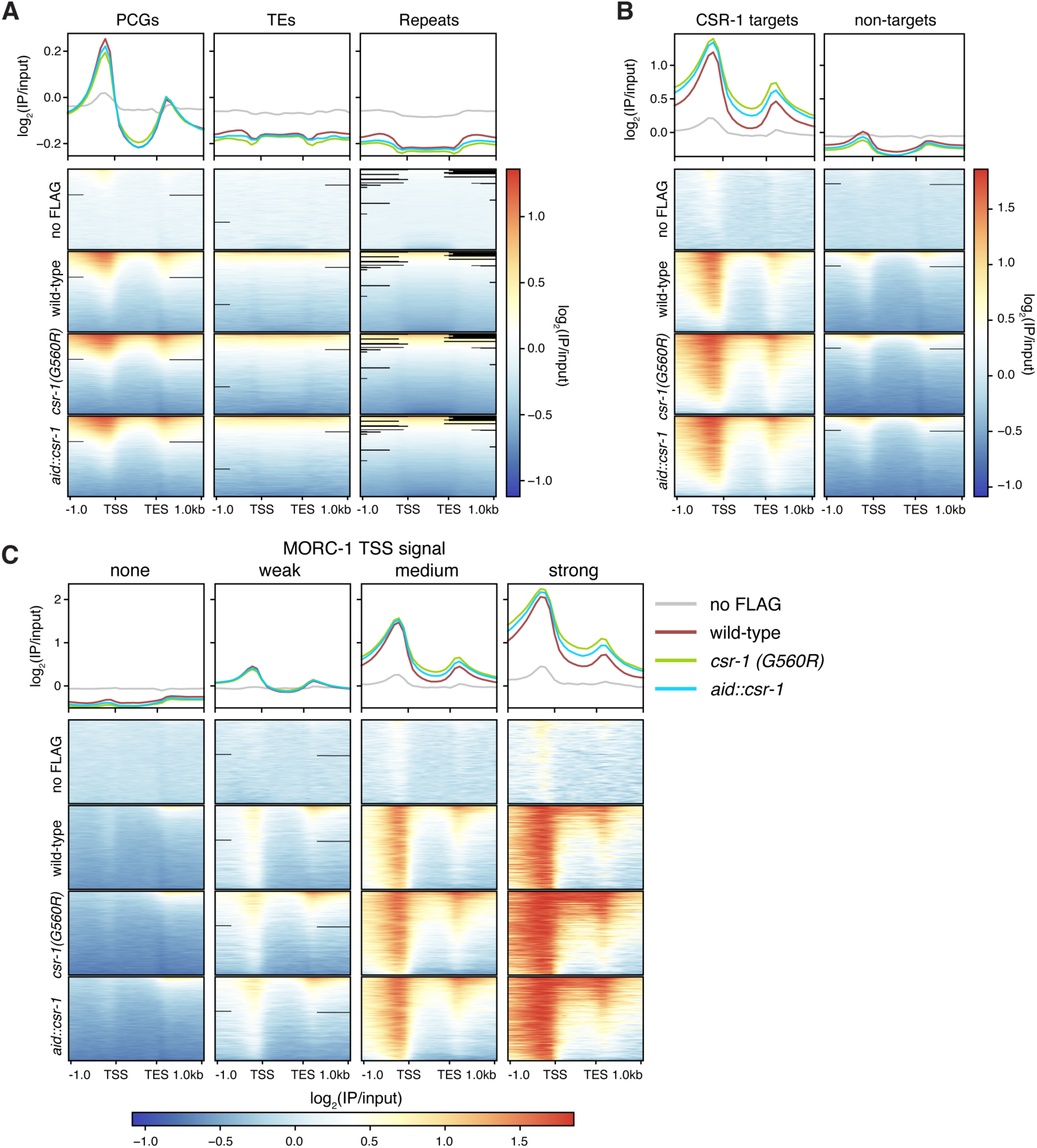
MORC-1 localization changes when overexpressed. (**A**) Metaplots and heatmaps showing MORC-1::FLAG localization (log_2_(IP/input)) in wild-type and both *csr-1(G560R)* and *aid::csr-1* [(+) auxin], over protein coding genes (PCGs), transposons (TEs), and repeat regions. No-FLAG control sample (wild-type worms lacking the 3xFLAG) shown for reference. Legend same as C. (**B**) Metaplots and heatmaps showing MORC-1::FLAG localization in wild-type and both *csr-1(G560R)* and *aid::csr-1* [(+) auxin], over CSR-1 target genes vs. non-targets. Legend same as C. (**C**) Metaplots and heatmaps showing MORC-1 localization in wild-type and both *csr-1(G560R)* and *aid::csr-1* [(+) auxin], over genes divided by wild-type MORC-1 occupancy at TSS.

**Fig. S8.**
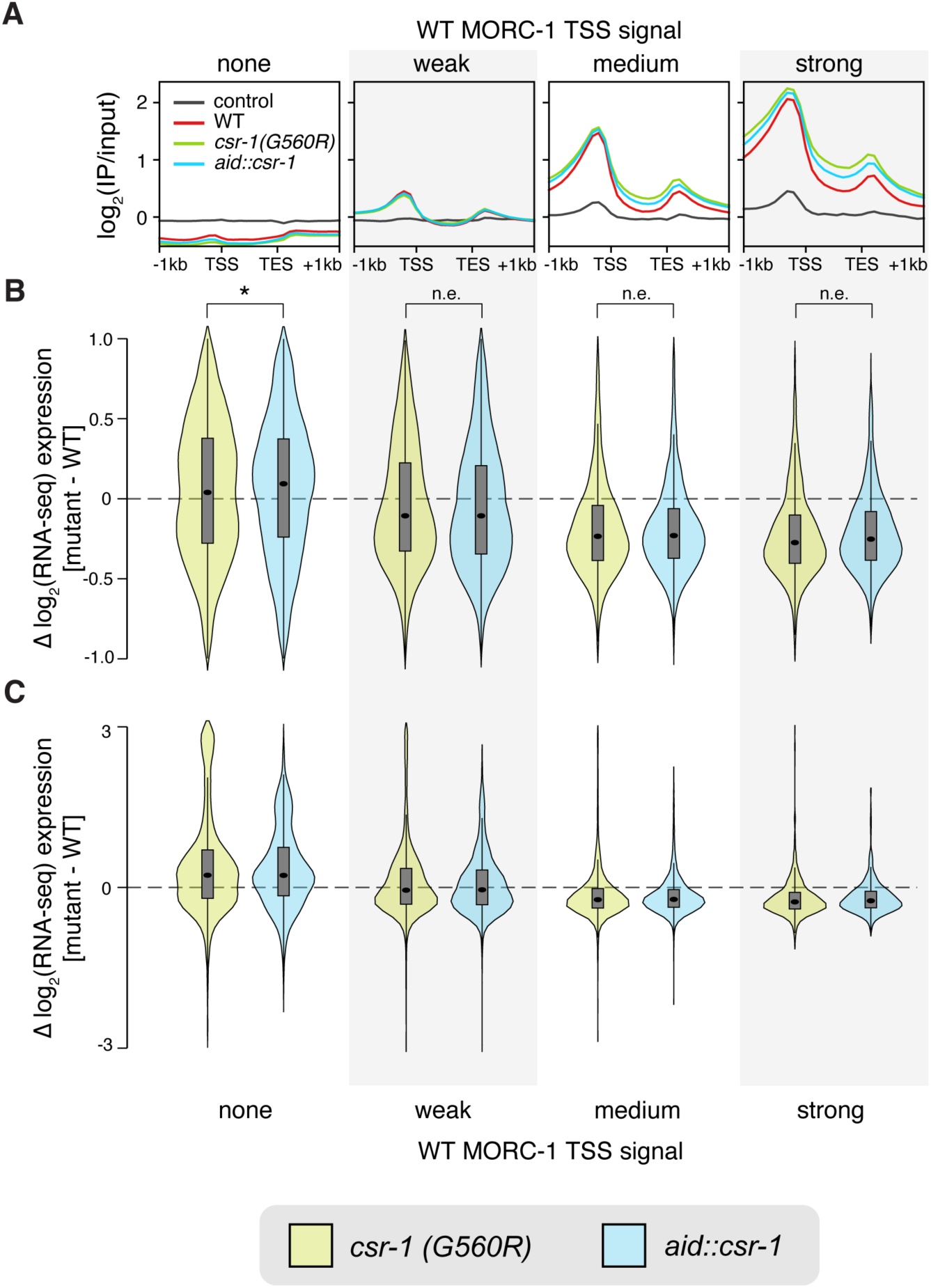
Consistent downregulation of genes highly bound by MORC-1 in both *csr1(G560R)* and *aid::csr-1*. (**A**) Metaplots of MORC-1 ChIP-seq signal in wild-type, *csr-1(G560R)* and *aid::csr-1* [(+) auxin] over genes binned by wild-type MORC-1 signal over TSS (same as Fig. 2D). (**B**) Distribution of log_2_ fold change expression values of indicated mutant over wild-type, estimated by DESeq2 (*35*), across genes binned by wild-type MORC-1 level at TSS as in (A). A small number of genes outside of y in [-1,1] not shown. (**C**) Same as (B), but expanding y-axis to show all values within [-3,3]. This also shows a population of genes weakly expressed in WT (low MORC-1) that become strongly upregulated in both mutants (see fig. S19, supplemental text). Effect size measured using Cohen’s d. n.e. = no/minimal effect (|d| < 0.2), * = |d| > 0.2, ** = |d| > 0.5, *** = |d| > 0.9, **** = |d| > 1.5.

**Fig. S9.**
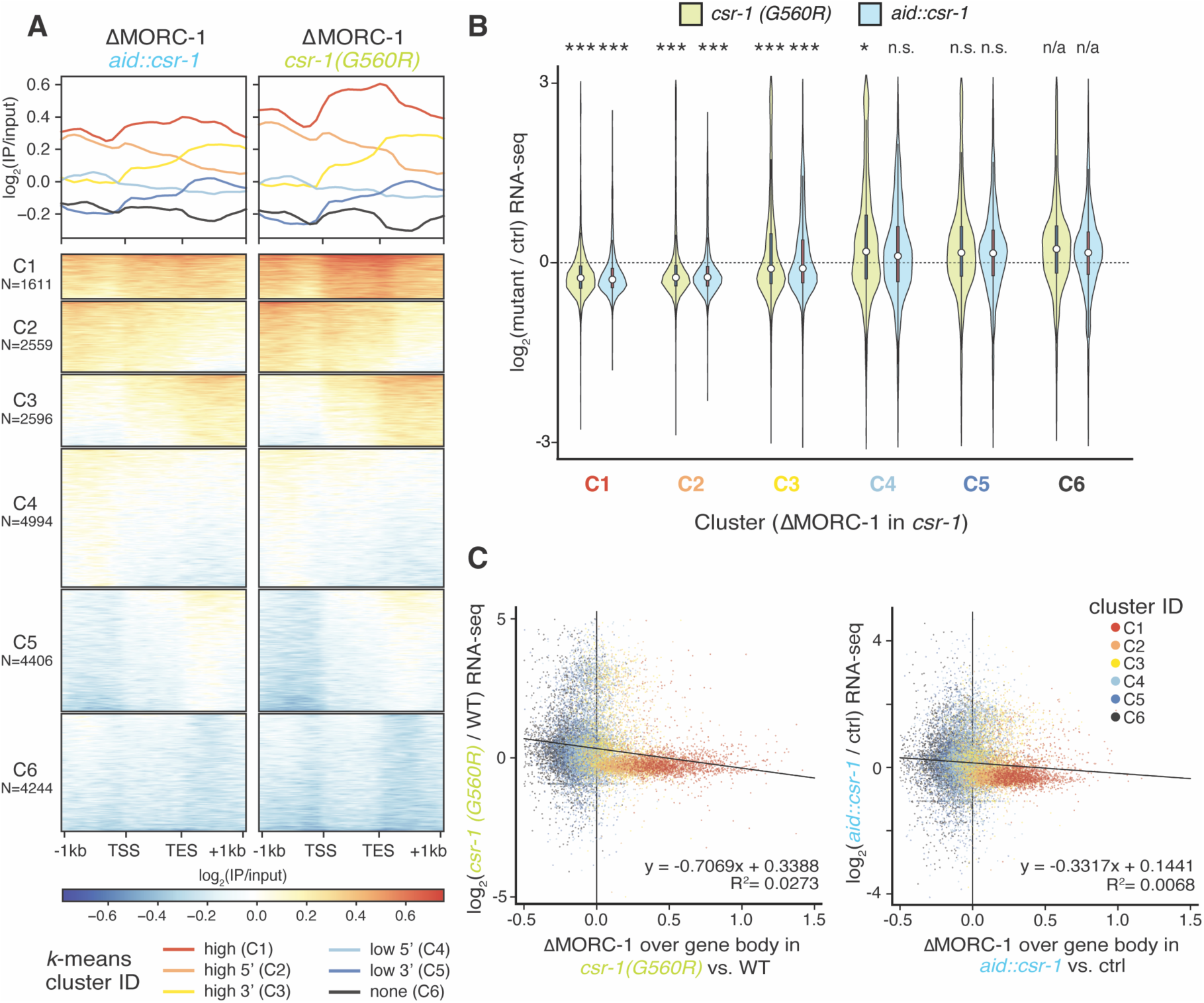
Correlation between MORC-1 gain in *csr-1* and gene downregulation. (**A**) Metaplots and heatmaps of change in MORC-1 levels in both *csr-1(G560R)* and *aid::csr-1* [(+) auxin] vs. control (difference in log_2_(IP/input) values) over all protein-coding genes. Genes were clustered using the *k*-means algorithm into six clusters, named C1-C6. Highest MORC-1 gain in *csr-1* occurs over genes in C1, while MORC-1 is lost at genes in C6. (**B**) Distribution of log_2_ fold change expression values of indicated mutant over control, estimated by DESeq2, across gene clusters from (A). A small number of genes with y not within [-3,3] not shown. Significance testing: *** = p < 0.0001, ** = p < 0.001, * = p < 0.01, n.s. = not significant; two-sample Wilcoxon rank-sum test with the null hypothesis that the observed distribution of log_2_ fold change values is drawn from the same distribution as cluster 6 genes (C6, rightmost violin plots). (**C**) Scatterplots of change in MORC-1 levels over the gene body in *csr-1(G560R)* (left panel) or *aid::csr-1* [(+) auxin] (right panel) compared to appropriate control, vs. the change in RNA-seq expression in the same mutant vs. control (DESeq2 estimated log_2_(fold change)). Genes are colored according to their cluster in (A). Black line indicates linear regression best fit line, equation and R^2^ indicated at bottom right. (A-C) For *csr-1(G560R)*, the control is N2 while for *aid::csr-1* all comparisons are between *aid::csr-1* worms treated with 1 mM auxin [(+) auxin] vs. 0 mM auxin [(-) auxin].

**Fig. S10.**
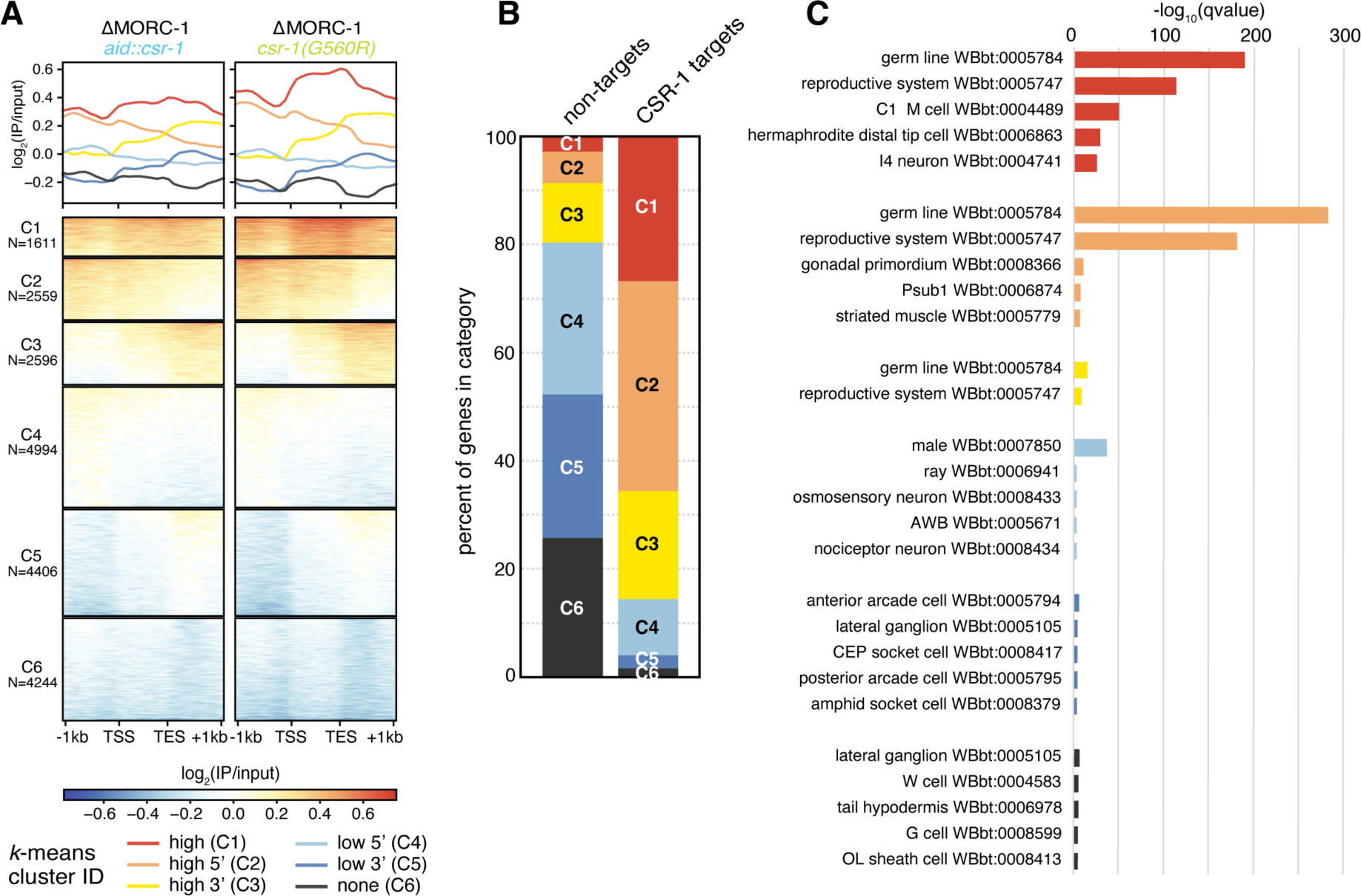
Characterization of MORC-1 gain clusters. (**A**) Same as fig. S9A. Metaplots and heatmaps of change in MORC-1 levels in both *csr-1(G560R)* and *aid::csr-1* [(+) auxin] vs. control (difference in log2(IP/input) values) over all protein-coding genes. Genes were clustered using the k-means algorithm into six clusters, named C1-C6, where C1 genes show strongest gain of MORC-1 in *csr-1*. (**B**) Percent of genes in each cluster from (A) that are CSR-1 targets vs. non-targets. (**C**) Tissue enrichment analysis using the WormBase Enrichment Analysis tool (*56*) for genes in each cluster from (A).

**Fig. S11.**
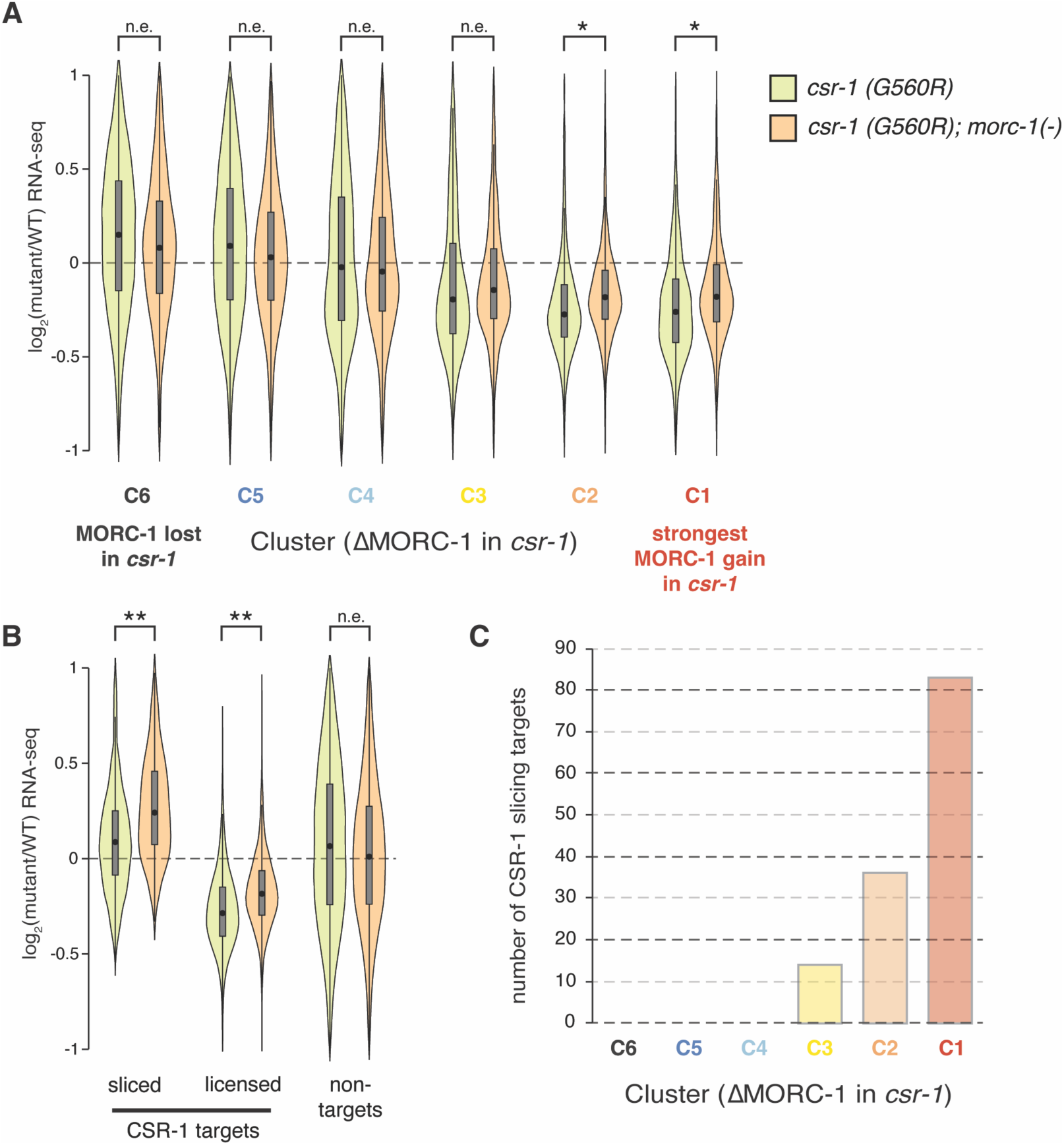
Target gene expression defects in *csr-1* are rescued by *morc-1* at genes highly bound by MORC-1. (**A-B**) Distribution of log_2_ fold change expression values of indicated mutant over wild-type, estimated by DESeq2, over (A) genes clustered based on change in MORC-1 levels in both *csr-1* mutants vs. WT, using k-means clustering with k == 6 (fig. S9A), and (B) CSR-1 targets vs. non-targets. Stars indicate effect size measured by Cohen’s d: n.e. = no/minimal effect (|d| < 0.1), * = |d| > 0.1, ** = |d| > 0.25, *** = |d| > 0.5, **** = |d| > 1. (C) Number of CSR-1 slicing targets (N=133, Gerson-Gurwitz *et al.* 2016 (*10*)) that fall into each of the 6 *k-*means clusters based on change in MORC-1 levels in *csr-*1 vs. WT (fig. S9A).

**Fig. S12.**
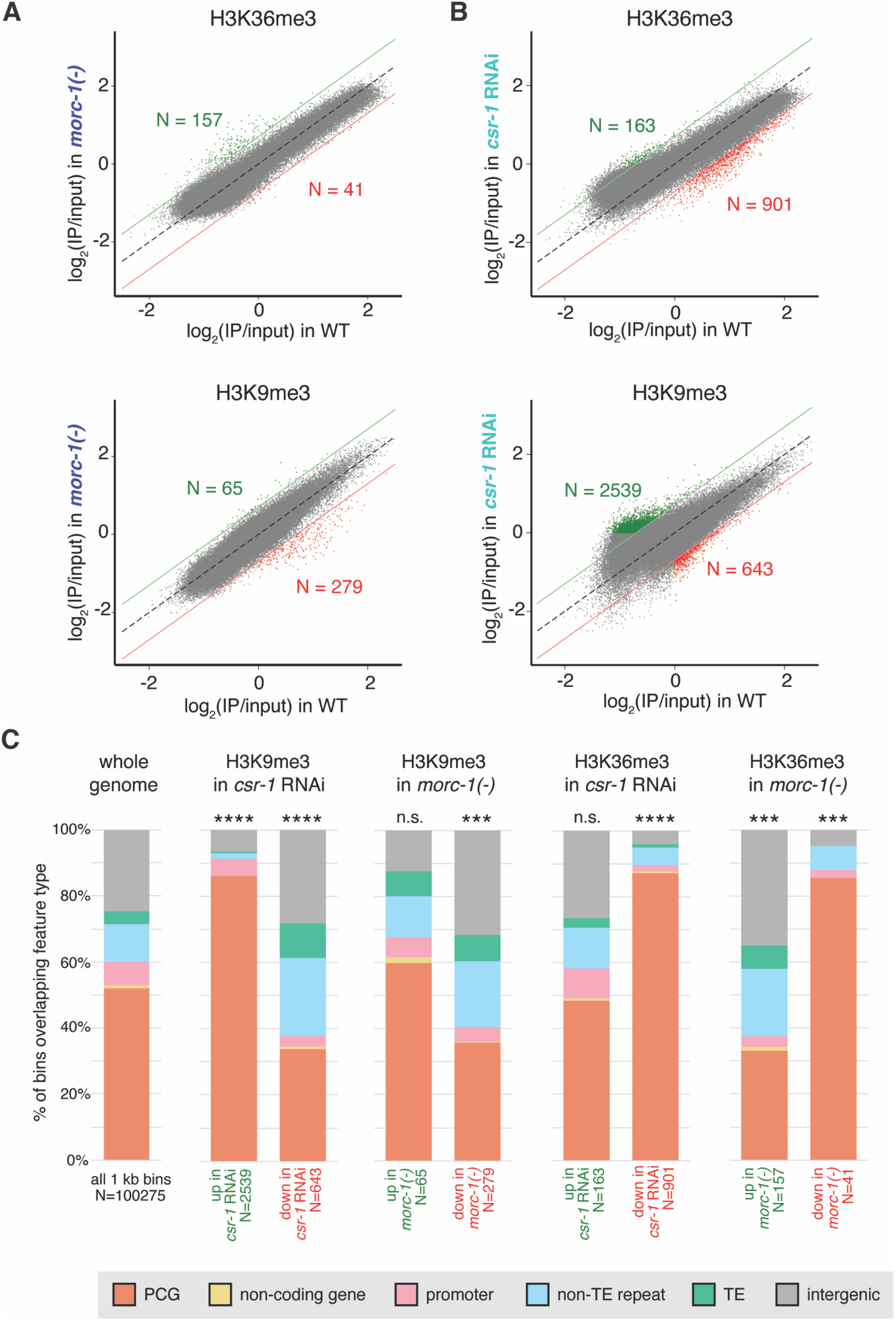
Genome-wide chromatin changes in *morc-1*(*-*) and *csr-1* RNAi. (**A-B**) Scatterplots of average H3K36me3 and H3K9me3 ChIP-seq signal (log_2_(IP/input)) in 1 kb bins tiled genome-wide. Wild-type (N2 + empty vector (EV) RNAi) signal is plotted against (A) *morc-1*(*-*) + EV RNAi or (B) N2 + *csr-1* RNAi. Significantly different bins, identified as +/-0.7 difference in log_2_(IP/input) between wild-type and *morc-1*(*-*) or *csr-1* RNAi, are highlighted in green and red (black dotted line: y = x line, green line: y = x + 0.7, red line: y = x – 0.7). (**C**) Distribution of genomic features overlapped by the 1 kb bins identified in A-B as having increased, decreased, or remained unchanged H3K9me3 or H3K36me3 in *csr-1* RNAi or *morc-1*(*-*). Significant deviation from the genome-wide distribution (leftmost panel) was evaluated using a Chi-squared test. Significance testing: (A-B) **** = p < 0.0001, *** = p < 0.001, ** = p < 0.01, * = p < 0.05, n.s. = not significant, two-tailed t-test.

**Fig. S13.**
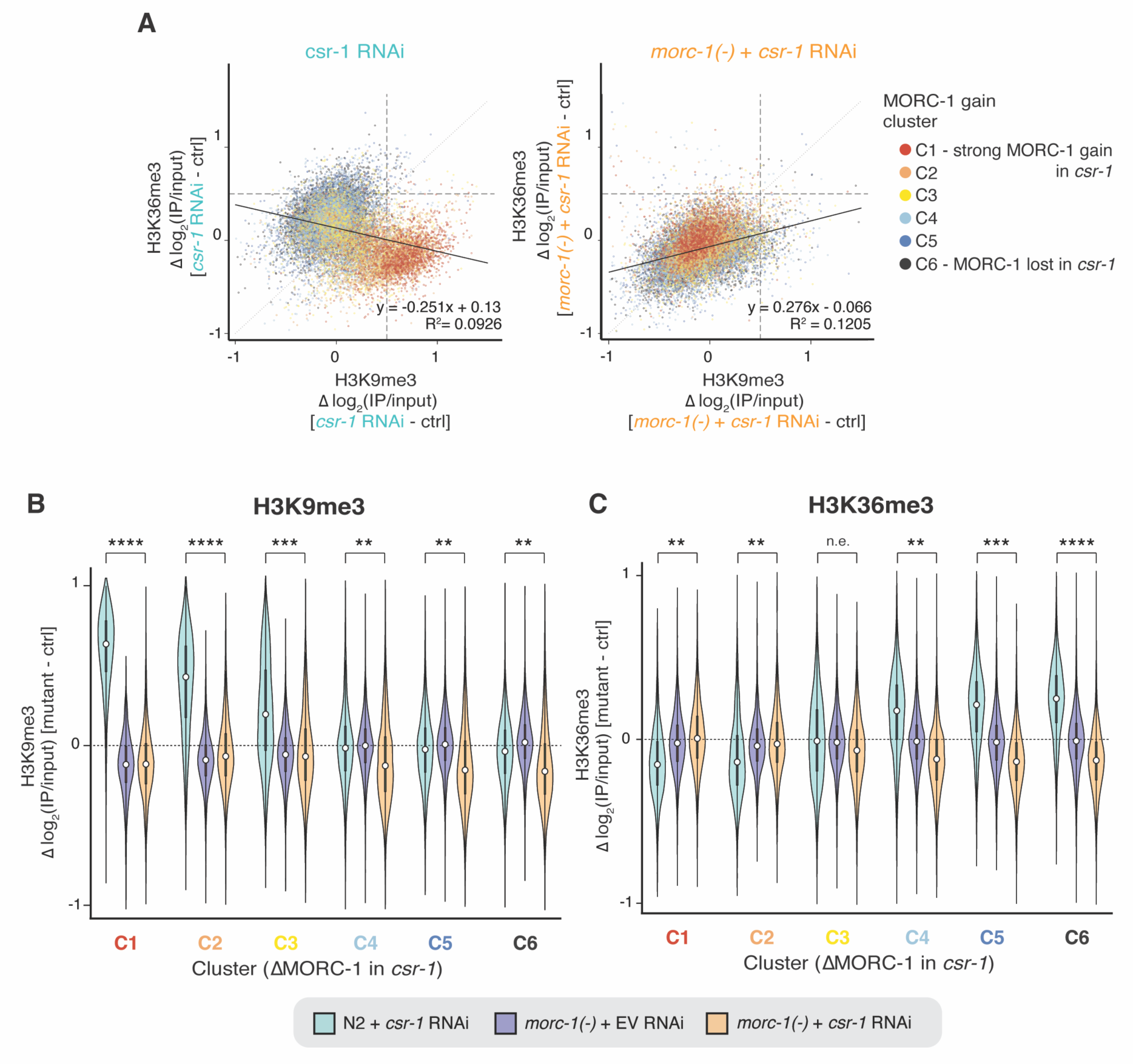
Changes in H3K9me3 and H3K9me3 in *csr-1* RNAi correlate with each other and are rescued by *morc-1*(*-*). (**A**) Scatterplots of change in H3K9me3 vs. change in H3K36me3 over gene bodies, in wild-type (N2) treated with *csr-1* RNAi or *morc-1*(*-*) treated with *csr-1* RNAi vs. control (N2 treated with empty vector (EV) RNAi). Value plotted is the difference in log_2_(IP/input) in indicated mutant/condition vs. control, so a value of 0 indicates that both the mutant and control had similar H3K9me3 or H3K36me3 values. Points are colored based on that gene’s *k*-means cluster from fig. S9A, with C1 (red) genes showing the highest MORC-1 gain in *csr-1*. Black line shows linear regression line, with equation and R^2^ at bottom right. (**B-C**) Distribution of change in (B) H3K9me3 and (C) H3K36me3 in indicated genotype + RNAi vs. control (in all cases, control is N2 + EV RNAi), over genes clustered based on change in MORC-1 levels in *csr-1* vs. WT (fig. S9A). Cohen’s d. n.e. = no/minimal effect (|d| < 0.2), * = |d| > 0.2, ** = |d| > 0.5, *** = |d| > 0.9, **** = |d| > 1.5.

**Fig. S14.**
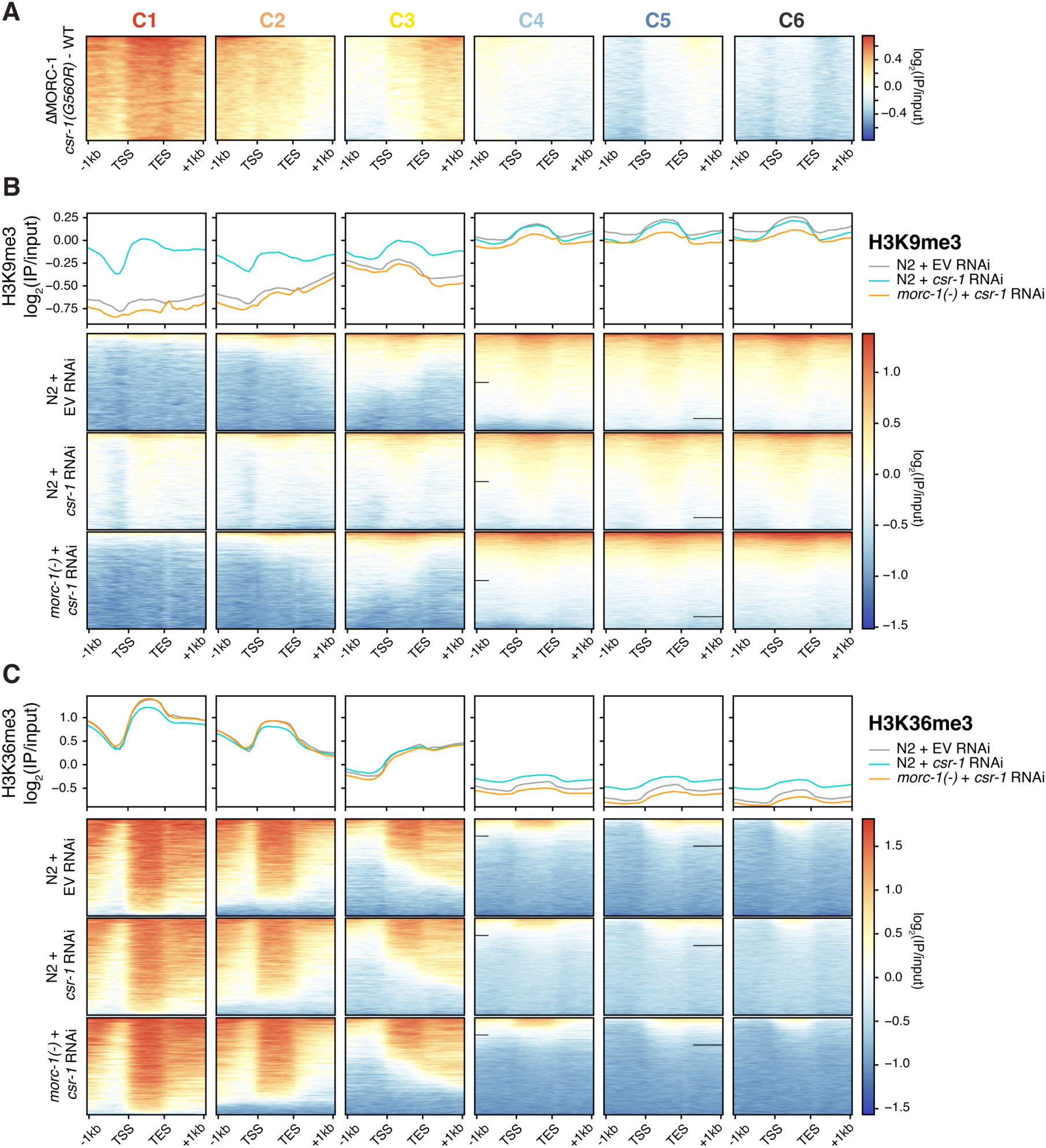
Changes in chromatin environment as a function of MORC-1 gain in *csr-1*(*-*). (**A**) Reproduction of heatmaps from fig. S9A, showing pattern of MORC-1 gain in each of the 6 *k-* means clusters in *csr-1(G560R)*. (**B-C**) Metaplots of (B) H3K9me3 and (C) H3K36me3, over genes in the 6 *k*-means clusters from fig. S9A, in wild-type worms treated with empty vector (EV) RNAi or *csr-1* RNAi, vs. *morc-1* worms treated with *csr-1* RNAi.

**Fig. S15.**
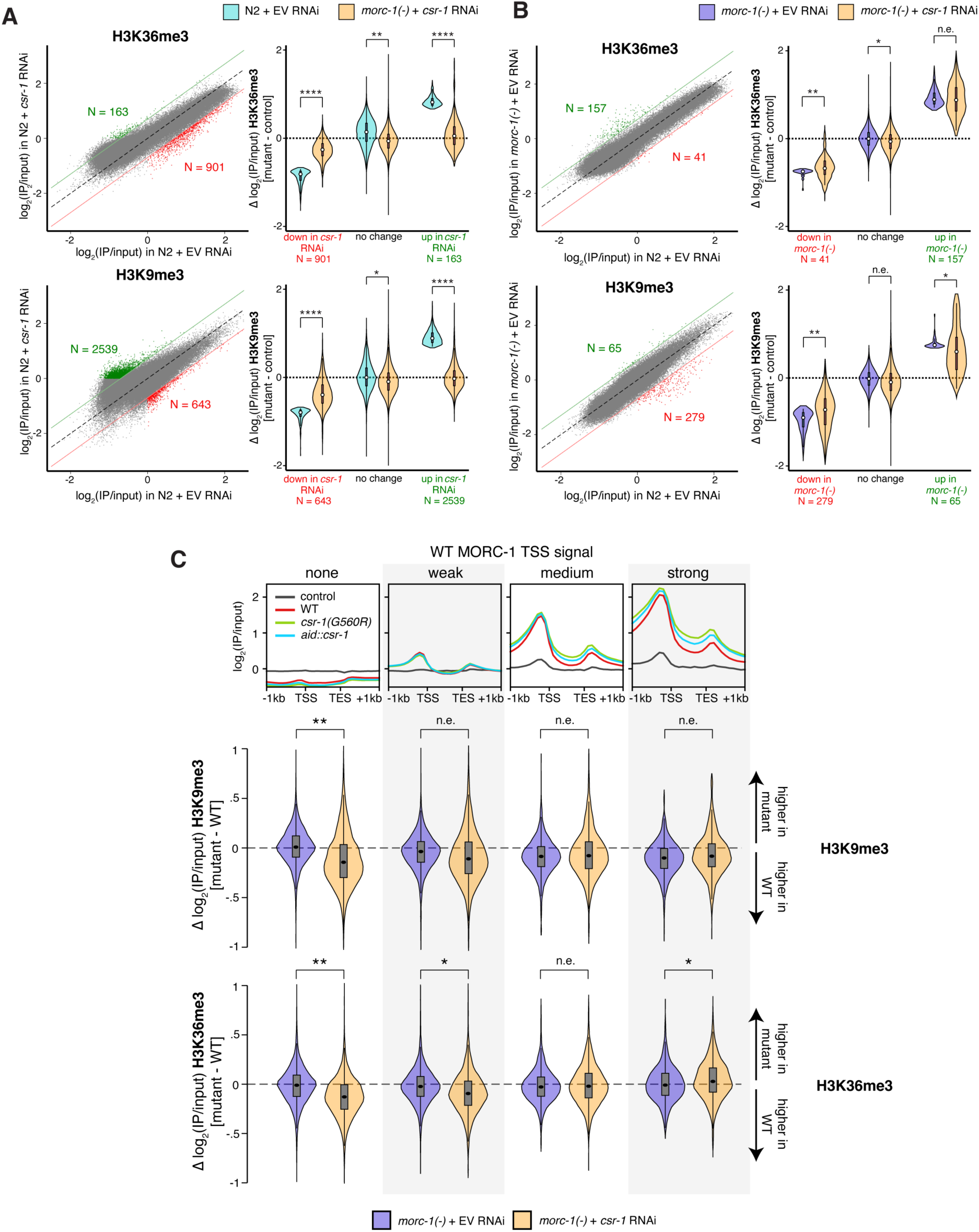
*morc-1*(*-*) can rescue chromatin defects in *csr-1* RNAi but not vice-versa. (**A-B**) Scatterplots same as fig. S12. For each scatterplot, the distribution of the difference in log_2_(IP/input) ChIP-seq signal between indicated genotype + RNAi and N2 + empty vector (EV) RNAi (WT) is plotted across up, down, and unchanged bins in each scatterplot as a violin plot, where y = 0 represents no difference from the control sample (N2 + EV RNAi). (A) demonstrates nearly complete rescue of *csr-1* RNAi changes by *morc-1*(*-*), while (B) demonstrates lack of rescue of *morc-1*(*-*) phenotype by *csr-1* RNAi. Significance testing: **** = p < 0.0001, *** = p < 0.001, ** = p < 0.01, * = p < 0.05, n.s. = not significant, two-tailed t-test comparing indicated distributions. (**C**) (Top) same as Fig. 2D: metaplot of log_2_(IP/input) ChIP-seq signal for MORC-1::FLAG in wild-type germline, *csr-1(G560R)* and *aid::csr-1*, over genes binned by wild-type MORC-1 levels at the TSS. (Bottom) Distribution of change in H3K9me3 and H3K36me3 in *morc-1*(*-*) treated with EV RNAi or *csr-1* RNAi, vs. N2 treated with EV RNAi, over genes binned by wild-type MORC-1 levels at the TSS (same as top). Effect size measured using Cohen’s d. n.e. = no/minimal effect (|d| < 0.2), * = |d| > 0.2, ** = |d| > 0.5, *** = |d| > 0.9, **** = |d| > 1.5. Note that these comparisons give p ∼ 0 by Student’s t-test or similar, due to large sample size.

**Fig. S16.**
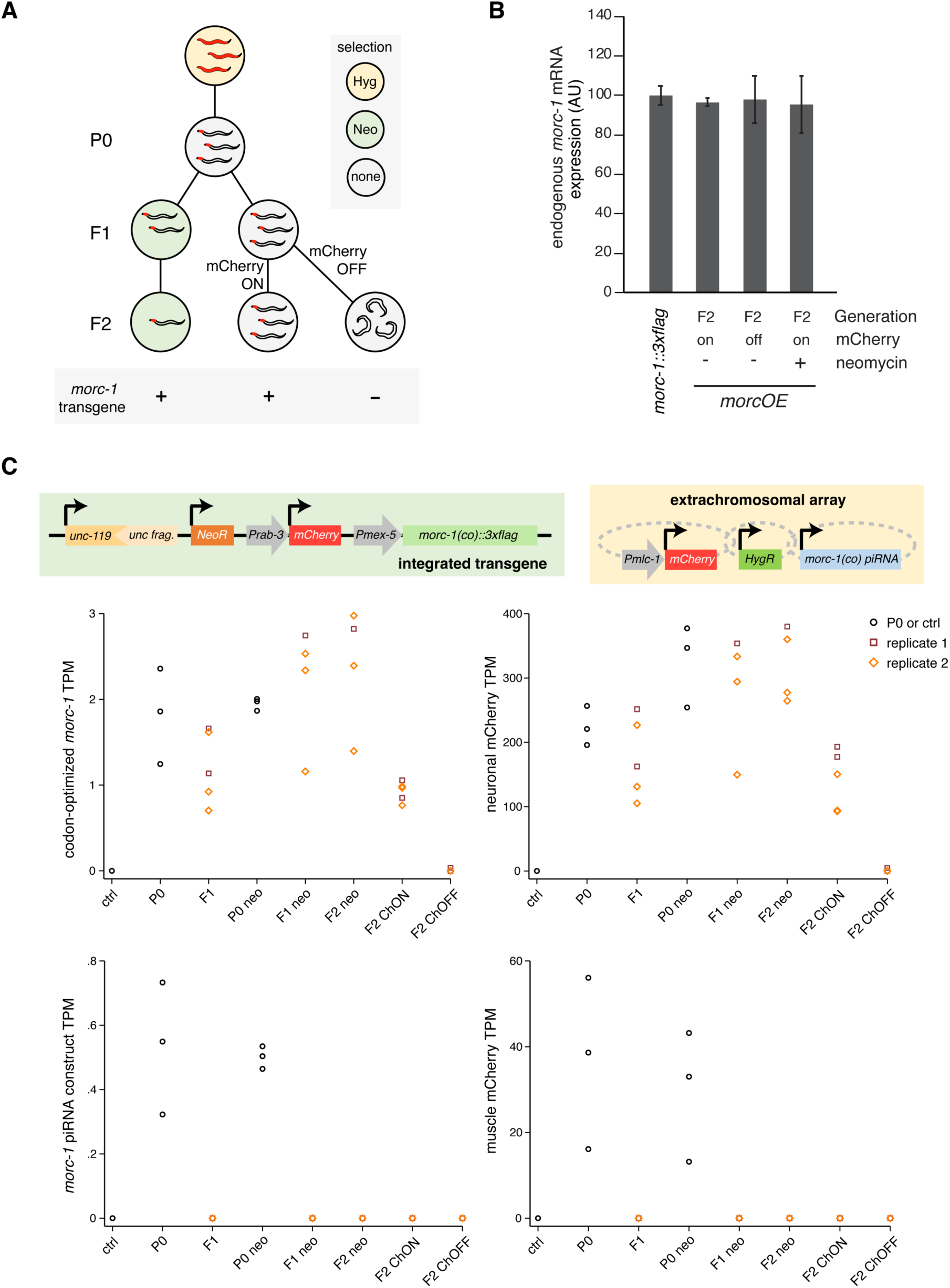
Expression of genes on integrated *morc-1* transgene and extrachromosomal array in *morcOE* samples. (**A**) Schematic of experimental design (same as Fig. 3D). (**B**) mRNA qPCR for the endogenous *morc-1* transcript in F2 *morcOE* worms vs. a non-transgenic control (MORC-1::3xFlag CRISPR strain). Error bars represent standard deviation of two technical replicates (**C**) TPM estimates from each RNA-seq replicate for codon-optimized MORC-1 and neuronal mCherry (both on integrated transgene), and the piRNAi construct and muscle mCherry (both on extrachromosomal array). P0 worms were split into two sets of replicates, which were taken through the experiment in parallel and are labelled here as ‘replicate 1’ and ‘replicate 2’. These are considered biological replicates in all analyses. Diagram of transgene and extrachromosomal array at top for reference. Control is same as (B), an endogenous *morc-1::3xflag* strain that lacks the integrated *morcOE* transgene and piRNAi extrachromosomal array. ChON = neuronal mCherry detected, ChOFF = neuronal mCherry not detected (see Fig. 3B, methods).

**Fig. S17.**
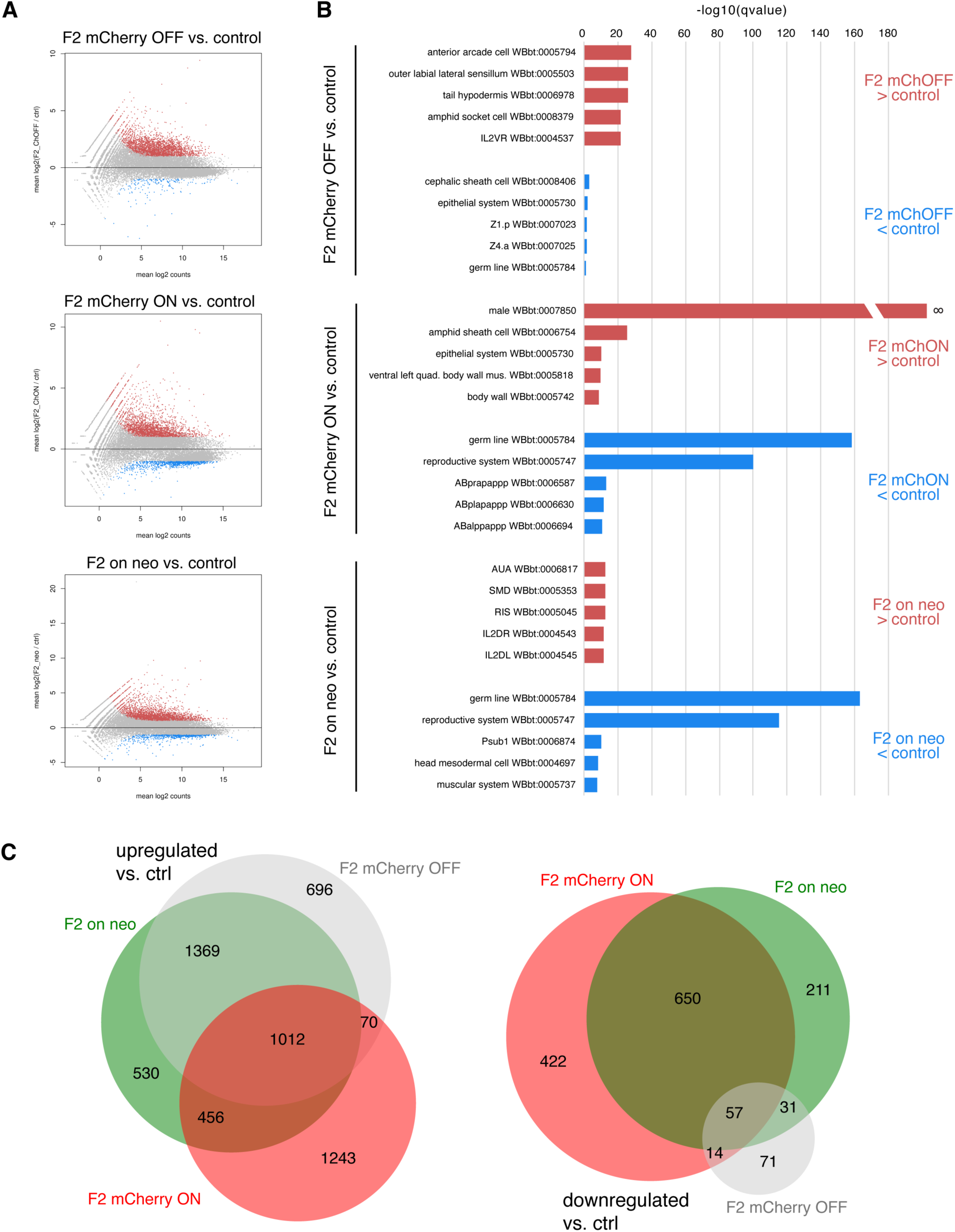
Characterization of genes up and downregulated in *morcOE* worms. (**A**) DESeq2 (*35*) MA plots for F2 worms expressing the transgene (F2 mCherry ON and F2 on neo) or those that have silenced the transgene (F2 mCherry OFF) vs. non-transgenic control (endogenous *morc-1::3xflag* only). (**B**) Q-values from tissue enrichment analysis (WormBase, see methods) for genes upregulated or downregulated in each group vs. the non-transgenic control. One term (F2 mCherry ON > ctrl, enrichment term “male”) had a q-value of 0, so the corresponding log10(qval) is infinite; here it is depicted as a broken bar passing beyond the y-axis of the plot. (**C**) Overlap between genes upregulated in each condition (left) and downregulated in each condition (right) compared to wild-type.

**Fig. S18.**
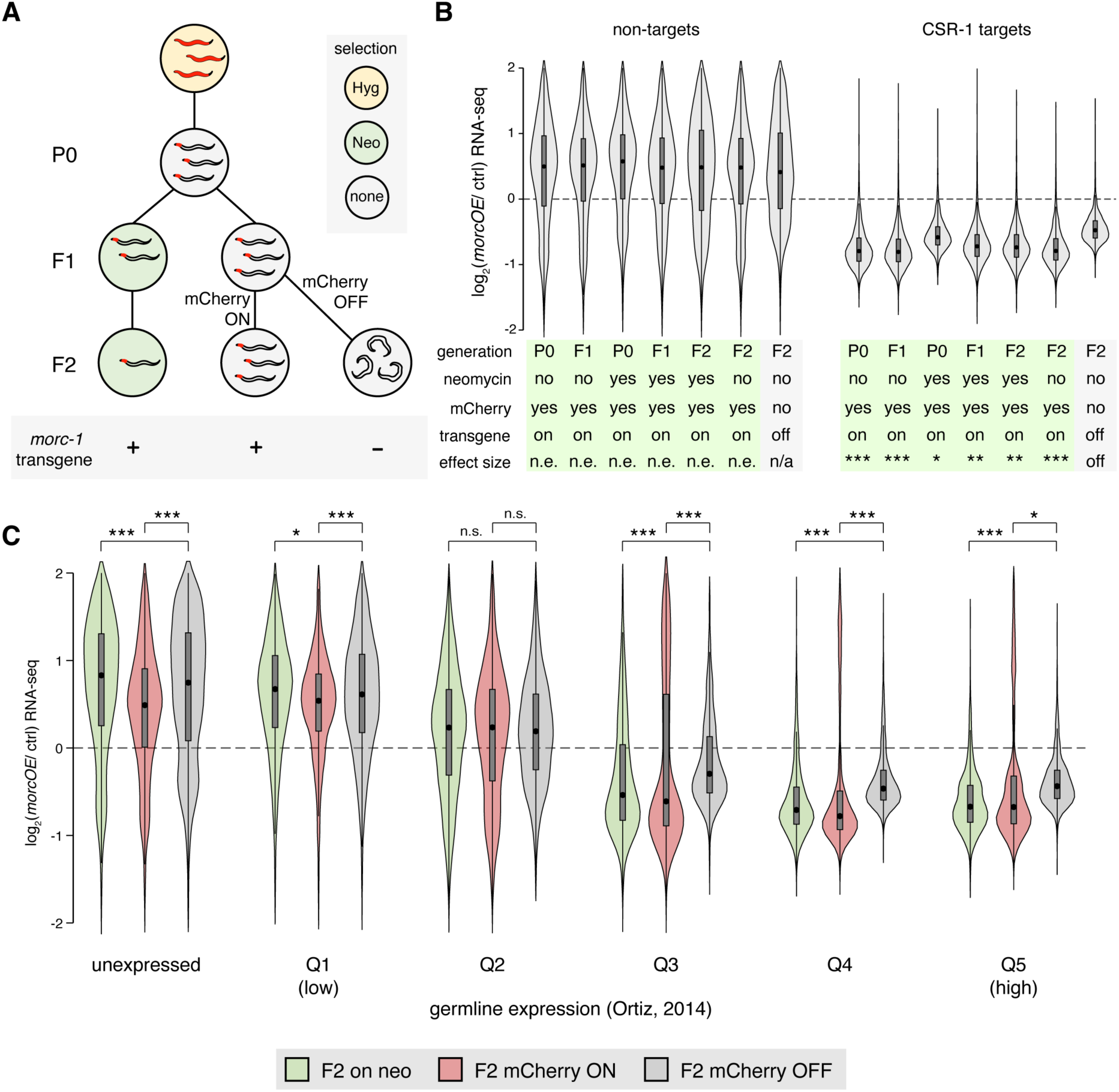
MORC-1 overexpression in wild-type germline downregulates germline-expressed genes and CSR-1 targets. (**A**) Diagram of experimental design used to separate worms that silenced the transgene from worms expressing it, same as Fig. 3D. (**B**) Distribution of expression changes in indicated sample, across CSR-1 targets (right) and non-targets (left). Bottom row indicates effect size compared to the F2 mCherry OFF sample (grey, right), measured by Cohen’s D, n.e. = no/minimal effect (|d| < 0.2), * = |d| > 0.2, ** = |d| > 0.5, *** = |d| > 0.9, **** = |d| > 1.5. (**C**) Distribution of expression changes in F2 samples that express the integrated *morc-1* transgene (F2 on neo and F2 mCherry ON) vs. those that have silenced the transgene (F2 mCherry OFF), across genes binned according to their expression in the germline in Ortiz *et al*. 2014 (*23*). n.s. = not significant, * p < 0.01, ** p < 0.001, *** p < 0.0001, two-tailed t-test.

**Fig. S19.**
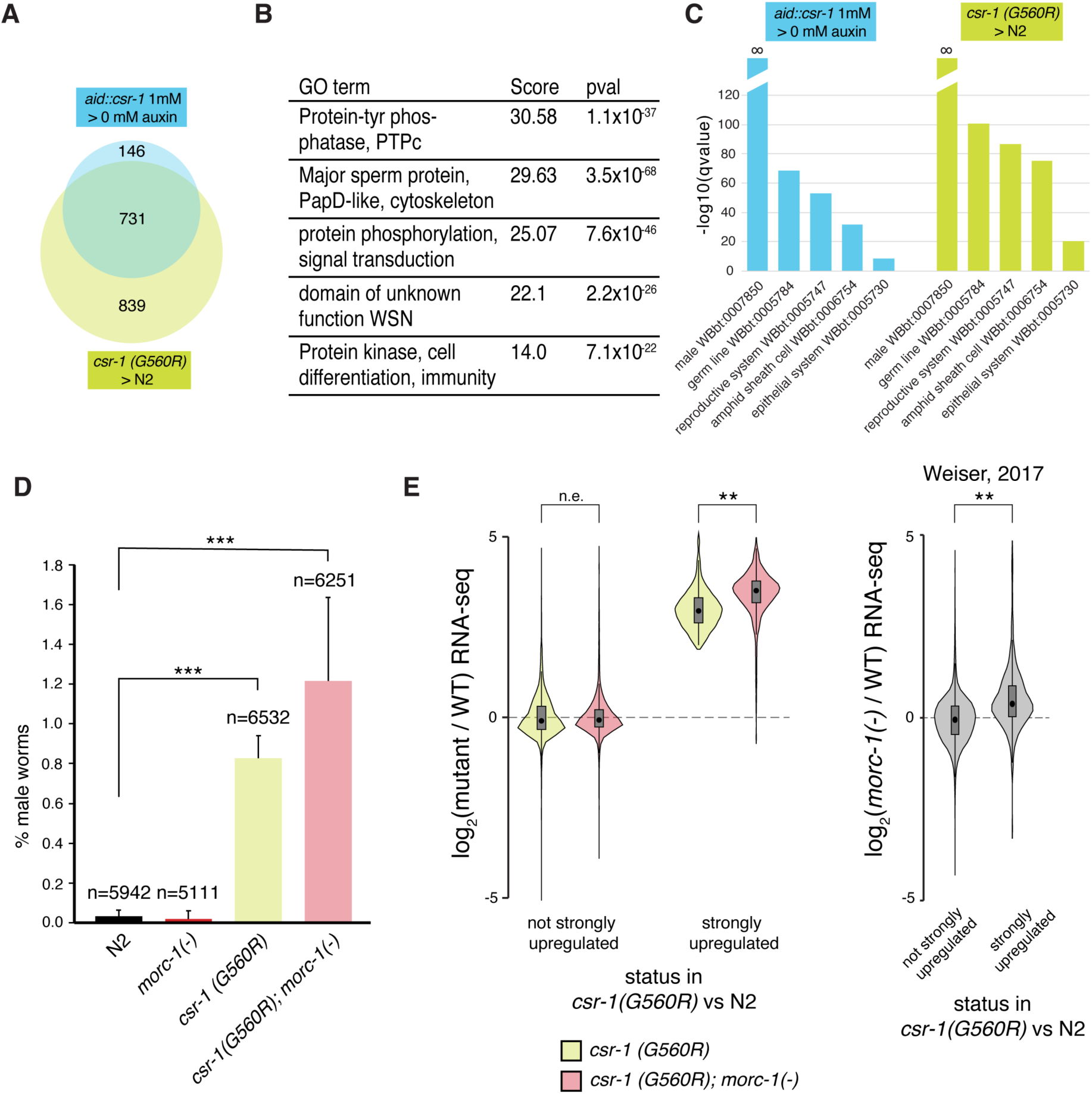
*him* phenotype in *csr-1(G560R)* causes upregulation of sperm-related genes and is not rescued by *morc-1*(*-*). (**A**) Overlap between genes significantly upregulated in *csr-1 (G560R)* vs. those significantly upregulated in *aid::csr-1* (log_2_ fold change > 1 & adjusted p-value < 0.01). (**B**) Top 5 GO-terms for genes significantly upregulated in *csr-1 (G560R)* vs. WT (log_2_ fold change > 1 & adjusted p-value < 0.01) (DAVID (*55*), see methods). (**C**) Top 5 tissue terms enriched among genes significantly upregulated in *csr-1 (G560R)* or *aid::csr-1*, WormBase enrichment analysis. (**D**) % male worms among indicated number of progeny from wild-type (N2), *morc-1*(*-*), *csr-1(G560R)* and *csr-1(G560R); morc-1*(*-*) worms. Error bars = standard deviation from 3 independent experiments. *** = p < 0.001, t-test. (**E**) Expression changes in (left) *csr-1(G560R)* and *morc-1*(*-*)*; csr-1(G560R)* vs. WT (this study) and (right) *morc-1*(*-*) vs. WT (Weiser et al. 2017 (*15*)), over n=1258 genes strongly upregulated in *csr-1 (G560R)* compared to WT (log_2_ fold change > 2 & adjusted p-value < 0.01), showing that *morc-1*(*-*) mildly amplifies this upregulation in *csr-1(G560R)*. Effect size measured by Cohen’s D, n.e. = no/minimal effect (|d| < 0.2), * = |d| > 0.2, ** = |d| > 0.5, *** = |d| > 0.9, **** = |d| > 1.5.

**Table S1.**
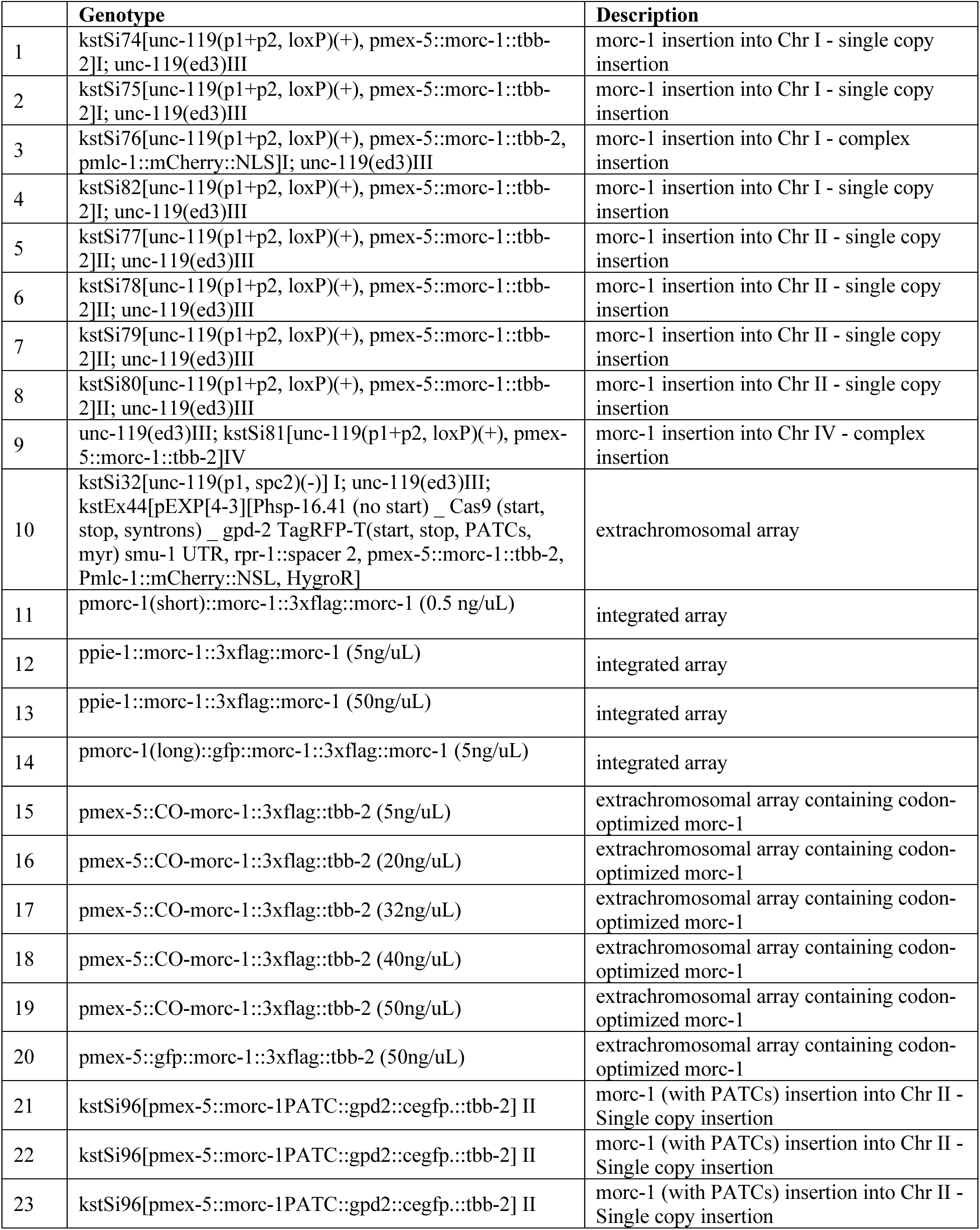
Failed attempts to generate a strain to overexpress *morc-1* in the germline, from which we were not able to recover worms overexpressing *morc-1* due to sterility.

**Table S2.** List of published sequencing datasets used in this study. N2 = wild-type strain Bristol N2.

|  | Seq Type | Genotype | ChIP | Tissue | SRA/Encode ID(s) | Ref |
| --- | --- | --- | --- | --- | --- | --- |
| 1 | ChIP-chip | N2 | H3K4me3 | whole worms | Encode dataset: 3552 | (59) |
| 2 | ChIP-seq | N2 | H3K27ac | germline | SRR9214969 | (22) |
| 3 | ChIP-seq | N2 | H3K27ac | germline | SRR9214971 | (22) |
| 4 | ChIP-seq | N2 | input | germline | SRR9214970 | (22) |
| 5 | ChIP-seq | N2 | input | germline | SRR9214972 | (22) |
| 6 | ChIP-seq | N2 | H3K36me3 | whole worms | SRR5024048 | (15) |
| 7 | ChIP-seq | N2 | H3K36me3 | whole worms | SRR5024049 | (15) |
| 8 | ChIP-seq | N2 | H3K36me3 | whole worms | SRR5024050 | (15) |
| 9 | ChIP-seq | *morc-1(-)* | input | whole worms | SRR5024036 | (15) |
| 10 | ChIP-seq | *morc-1(-)* | H3K36me3 | whole worms | SRR5024052 | (15) |
| 11 | ChIP-seq | *morc-1(-)* | H3K36me3 | whole worms | SRR5024053 | (15) |
| 12 | ChIP-seq | N2 | input | whole worms | SRR5024038 | (15) |
| 13 | ChIP-seq | *morc-1(-)* | H3K9me3 | whole worms | SRR5024028 | (15) |
| 14 | ChIP-seq | *morc-1(-)* | H3K9me3 | whole worms | SRR5024030 | (15) |
| 15 | ChIP-seq | N2 | H3K9me3 | whole worms | SRR5024031 | (15) |
| 16 | RNA-seq | N2 | n/a | whole worms | SRR5024017 | (15) |
| 17 | RNA-seq | N2 | n/a | whole worms | SRR5024018 | (15) |
| 18 | RNA-seq | *morc-1(-)* | n/a | whole worms | SRR5024021 | (15) |
| 19 | RNA-seq | *morc-1(-)* | n/a | whole worms | SRR5024022 | (15) |
| 20 | RNA-seq | *fog-2(-)* | n/a | whole worms | SRR1263137 SRR1263138<br>SRR1263139 SRR1263140 | (23) |
| 21 | RNA-seq | *fog-2(-)* | n/a | whole worms | SRR1263141 SRR1263142<br>SRR1263143 SRR1263144 | (23) |
| 22 | RNA-seq | *fog-2(-)* | n/a | whole worms | SRR1263145 SRR1263146<br>SRR1263147 SRR1263148 | (23) |
| 23 | RNA-seq | *fog-2(-)* | n/a | whole worms | SRR1263149 SRR1263150<br>SRR1263151 SRR1263152 | (23) |
| 24 | RNA-seq | *fog-2(-)* | n/a | whole worms | SRR1263153 SRR1263154<br>SRR1263155 SRR1263156 | (23) |
| 25 | RNA-seq | *fog-2(-)* | n/a | whole worms | SRR1263157 SRR1263158<br>SRR1263159 SRR1263160 | (23) |
| 26 | RNA-seq | *fog-2(-)* | n/a | whole worms | SRR1263161 SRR1263162<br>SRR1263163 SRR1263164 | (23) |
| 27 | RNA-seq | *fog-2(-)* | n/a | whole worms | SRR1263165 SRR1263166<br>SRR1263167 SRR1263168 | (23) |
| 28 | RNA-seq | *fem-3(-)* | n/a | whole worms | SRR1263169 SRR1263170<br>SRR1263171 SRR1263172 | (23) |
| 29 | RNA-seq | *fem-3(-)* | n/a | whole worms | SRR1263173 SRR1263174<br>SRR1263175 SRR1263176 | (23) |
| 30 | RNA-seq | *fem-3(-)* | n/a | whole worms | SRR1263177 SRR1263178<br>SRR1263179 SRR1263180 | (23) |
| 31 | RNA-seq | *fem-3(-)* | n/a | whole worms | SRR1263181 SRR1263182<br>SRR1263183 SRR1263184 | (23) |
| 32 | RNA-seq | *fem-3(-)* | n/a | whole worms | SRR1263185 SRR1263186<br>SRR1263187 SRR1263188 | (23) |
| 33 | RNA-seq | *fem-3(-)* | n/a | whole worms | SRR1263189 SRR1263190<br>SRR1263191 SRR1263192 | (23) |
| 34 | RNA-seq | *fem-3(-)* | n/a | whole worms | SRR1263193 SRR1263194<br>SRR1263195 SRR1263196 | (23) |
| 35 | RNA-seq | *fem-3(-)* | n/a | whole worms | SRR1263197 SRR1263198<br>SRR1263199 SRR1263200 | (23) |

**Data S1. Mapping statistics for all sequencing data.** Number of reads that were obtained from sequencing, that passed QC filtering, that aligned uniquely, and that remained after PCR deduplication, for all libraries generated in this study, as well as all published sequencing datasets reanalyzed for this study using the same pipeline.

**Data S2. Full dataset used for analyses over genes.** Excel file containing three sheets. The first (“gene data”) contains per-gene level RNA-seq, sRNA-seq, ChIP-seq and ATAC-seq data used to generate most figures in this study. ChIP-seq data were averaged over either gene body or transcriptional start site (TSS) region (see methods). Also includes several other useful variables (flags for CSR-1 targets, published germline expression data, etc.). The second (“transgene data”) contains TPM estimates for all *morcOE* samples for expression of genes expressed either from the integrated transgene or from the extrachromosomal array in that line. The last sheet (“list of variables”) contains detailed information on each variable in both sheets.

**Data S3. H3K9me3 and H3K36me3 ChIP-seq signal over 1-kb bins genome-wide.** Average ChIP-seq signal (log2(IP/input)) across 1 kb bins tiled genome-wide. Includes H3K9me3 and H3K36me3 data (all replicates merged) for N2, N2 + *csr-1* RNAi, *morc-1*(*-*), and *morc-1*(*-*) + *csr-1* RNAi; used for figs. S12, S15. The “variable info” tab contains information on each variable.

**Data S4. Fertility & Him assay data.** Raw data for all fertility assays and high incidence of males (Him) assays. Fertility assays in F2 *morcOE* worms (Fig. 3C) compare our *morcOE* strain separated into transgene (tg) expressed or silent, vs. a non-transgenic control expressing only *morc-1::3xflag* from the endogenous locus. Status of each parent’s movement and mCherry phenotypes also recorded, as well as inferred *morc-1* transgene status (tg expressed or tg silent). Worms were transferred to new plates as progeny were laid to facilitate counting, and counts on each plate are recorded separately. This data was also collected across two different experiments, referred to as replicate 1 and 2. All other fertility assays (Fig. 1A, 1C, fig. S1B) are recorded in another tab, and only total progeny are listed. For Him assays, the experiment was repeated 3 times.

